# Analyzing the T cell receptor repertoire of 2,804 individuals with inflammatory bowel disease identifies public T cell responses involved in the pathogenesis

**DOI:** 10.1101/2025.07.24.666514

**Authors:** Hesham ElAbd, Aya K.H. Mahdy, Érika Endo Kokubun, Valeriia Kriukova, Christine Olbjørn, Gøri Perminow, May-Bente Bengtson, Petr Ricanek, Svend Andersen, Trond Espen Detlie, Vendel A. Kristensen, Johannes Roksund Hov, IBSEN-III study group, Marte Lie Høivik, Andre Franke

**Author notes:** Shared first co-authorship (Hesham ElAbd, Aya K.H. Mahdy).

## Abstract

Whereas altered immune processes have been identified in individuals with inflammatory bowel disease (IBD), potentially causative antigens remain to be identified. By interrogating the immune repertoire of individuals with IBD, an identification of common antigenic exposures associated with the disease can be obtained. We analyzed the T cell receptor beta (TRB) chain repertoire of 1,890 individuals with Crohn’s disease (CD) and 914 individuals with ulcerative colitis (UC), enabling the identification of 327 TRB clonotypes associated with CD and 130 with UC. We validated the expansion of these clonotypes in a cohort of treatment-naïve individuals with either CD, UC or symptomatic control (n=855). These disease-associated clonotypes were restricted to disease-associated risk HLA alleles and their expansion correlated with disease-severity but not with surgery or treatment trajectory. In conclusion, we identified and validated TRB clonotypes that are associated with either CD or UC, these clonotypes are a novel therapeutic target in IBD.

## Introduction

Inflammatory bowel disease (IBD) is a chronic, idiopathic, inflammatory disease of the gastrointestinal tract (GIT) with two main forms, Crohn’s disease (CD) and ulcerative colitis (UC). Interactions among environmental factors^*1*^, the microbiome^*2, 3*^, Epstein-Barr virus (EBV) infection (infectious mononucleosis^*4*^), and genetic predisposition^*5–7*^ have been shown to contribute to disease development. Besides variants at the *NOD2*^*8, 9*^ and *ATG16L*^*10*^ gene loci, both being strongly associated with CD, multiple human leukocyte antigens (HLA) alleles have been associated with IBD, *e.g.* HLA-DRB1*07:01^*6, 11*^ with CD, particularly ileal CD, and HLA-DRB1*15:01 with UC^*5*^. Aberrant immune processes have been observed in individuals with IBD, such as alterations in responses toward gut microbiota^*12*^, increased cytotoxic responses toward fungal antigens^*13*^, and an expansion of a subset of type II invariant natural killer T cells in CD patients^*14*^. In addition, individuals with CD have been shown to have an altered B cell receptor repertoire^*15*^, and an increased antibody-response towards bacterial flagellins^*16, 17*^.

The T cell receptor (TCR) repertoire comprises the collection of TCRs encoding the immunological exposure history of an individual^*18*^. A common method to study the T cell repertoire is through bulk repertoire sequencing (TCR-Seq), which identifies thousands of TCR antigenic binding chains, *e.g.,* the alpha (TRA) or the beta (TRB) chains present in a sample. These chains are generated using a somatic recombination process termed V(D)J recombination, producing a large array of TRAs and TRBs chains (*i.e.,* clonotypes), which enables T cells to recognize a diverse set of antigens presented by HLA proteins. While TCR-Seq does not provide the antigenic target of these TRAs or TRBs clonotypes, large-scale statistical analyses can be utilized to associate a particular clonotype with a specific trait. For example, by profiling the TRB repertoire of 666 healthy donors with known cytomegalovirus (CMV) serostatus, 164 clonotypes associated with CMV infection were identified^*19*^. The same framework was used to discover the public immune signature of SARS-CoV2 infection^*20*^, Lyme disease^*21*^ and type I diabetes^*22*^.

Recently, TCR-Seq has been used to decode the immune signature of IBD^*23*^, nonetheless, the identified clonotypes has not been validated in different cohorts. Furthermore, the cohort was mostly assembled from treated individuals and hence the discovered clonotypes can be a consequence of treatment and not the disease. Hence, to resolve this, we analyzed the TRB repertoire of the US-based “*A Study of a Prospective Adult Research Cohort with IBD (SPARC IBD)*” cohort of the Crohn’s & Colitis Foundation IBD Plexus research program^*24*^ to discover clonotypes that are either associated with CD or UC. Subsequently, we validated the expansion of these clonotypes in a cohort of treatment-naïve and treated individuals from Norway, showing that they are mainly expanded because of the disease and not the treatment.

## Results

### Identifying CD- and UC-associated clonotypes

By comparing the repertoire of individuals with CD to that of individuals with UC, we identified 327 clonotypes associated with CD (**Table S3**) and 130 clonotypes associated with UC (**Table S4**) (**Material and Methods**)^*19*^. The expansion of CD-associated clonotypes was higher in individuals with CD relative to individuals with UC (**Fig. 2A**). Conversely, the expansion of UC-associated clonotypes was higher in individuals with UC relative to individuals with CD (**Fig. 2B**). Utilizing seeded clustering (**Material and Methods**) the number of CD-associated clonotypes increased from 327 to 2,827 clonotypes, arranged into 327 meta-clonotypes, with a median of seven and a maximum of 38 clonotypes per meta-clonotype (**Fig. S1A**). The clustering also increased the number of UC-associated clonotypes to 908 clonotypes arranged into 130 meta-clonotypes with a median of five and a maximum of 33 clonotypes per meta-clonotype (**Fig. S1B**). The extended set of CD-associated meta-clonotypes and UC-associated meta-clonotypes were significantly expanded in individuals with CD (**Fig. 2C**) and individuals with UC (**Fig. 2D**), respectively. Thus, for the rest of the manuscript, we focused our analyses on these meta-clonotypes unless stated otherwise.

**Figure 1:**
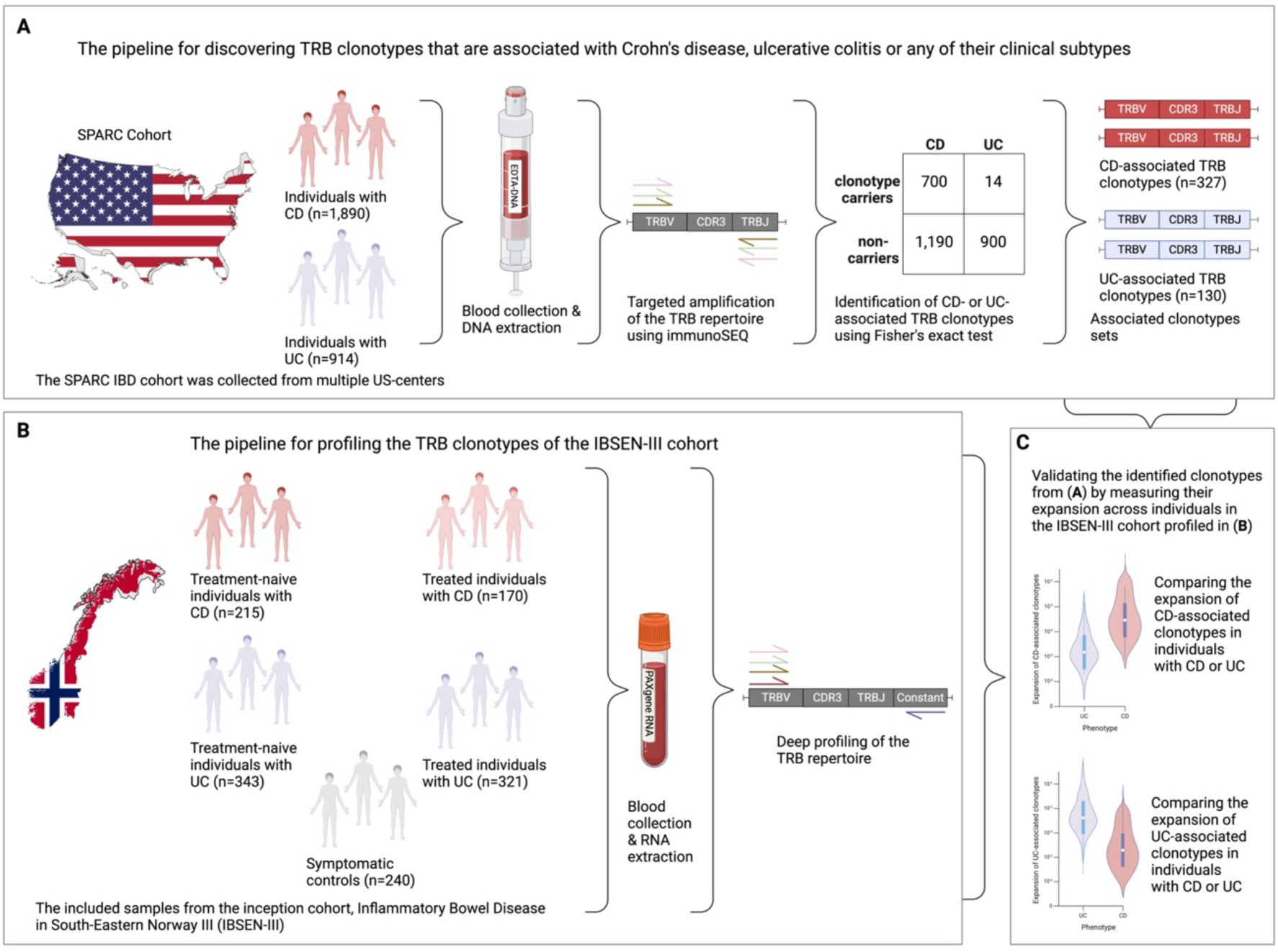
*Overview of the experimental approach used for the identification and validation of TRB clonotypes that are associated with CD or UC.* (**A**) illustrates the pipeline used for discovering TRB clonotypes that were associated with either CD or UC. 1,890 and 914 individuals with CD or UC, respectively, from the SPARC IBD cohort, were included in the study. These individuals were recruited from multiple centers across the US. After blood collection and DNA isolation, the TRB repertoire was profiled using the immunoSEQ assay. After cleaning and filtering the generated data, we used the incidence analysis framework proposed by Emerson et al.^*19*^ to identify clonotypes that are associated with either CD or UC. This enabled us to identify 327 TRB clonotypes that were associated with CD and 130 clonotypes that were associated with UC. (**B**) shows the composition of the IBSEN-III cohort and the workflow for profiling the TRB repertoire of the cohort. Specifically, we profiled the TRB repertoire of individuals with IBD and symptomatic controls at the time of diagnosis and before any treatment, i.e., treatment-naive samples and one year after treatment. From each individual, the TRB repertoire was profiled using RNA extracted from PAXgene tubes. (**C**) depicts the validation strategy of the clonotypes identified in (**A**) on the IBSEN-III cohort profiled in (**B**) by quantifying the expansion of CD- and UC-associated meta-clonotypes in treatment-naive and treated individuals with IBD.

**Figure 2:**
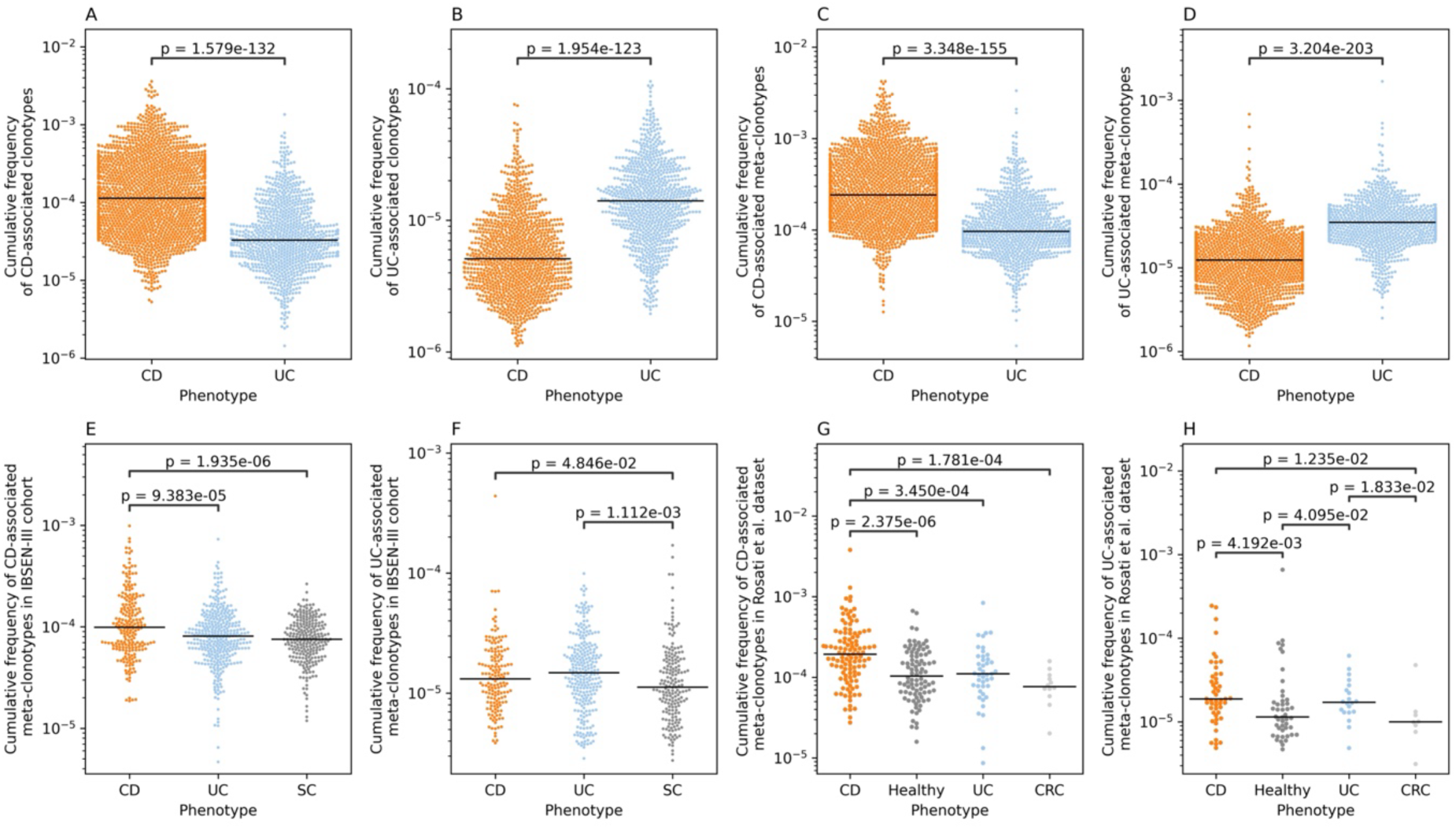
*The expansion of CD- and UC-associated clonotypes and meta-clonotypes in individuals from the SPARC IBD cohort and the two validation datasets, namely IBSEN-III and* from Rosati et al.^*14*^. (**A**) shows the expansion of CD-associated clonotypes in individuals with CD or UC from the SPARC IBD cohort, (**B**) the expansion of the UC-associated clonotypes in the same individuals. (**C**) depicts the expansion of CD-associated meta-clonotypes and (**D**) the expansion of UC-associated meta-clonotypes in individuals with CD or UC included in the SPARC IBD cohort. (**E**) shows the expansion of CD-associated meta-clonotypes in symptomatic controls (SC) vs. treatment-naive individuals with CD or UC from the IBSEN-III cohort, while (**F**) shows the expansion of UC-associated meta-clonotypes in the same individuals. (**G**) illustrates the expansion of CD-associated meta-clonotypes, and (**H**) shows the expansion of UC-associated meta-clonotypes in the blood TRB repertoire of individuals with CD, UC, or colorectal carcinoma (CRC) as well healthy controls in the test dataset derived from Rosati et al.^*14*^. Across all panels and figures – and unless stated otherwise – group comparisons were conducted using the two-sided Mann-Whitney U test, with only statistically significant comparisons shown. All panels show individuals where these clonotypes have been detected, i.e., individuals where the expansion of the meta-clonotypes was not detected are not shown.

Subsequently, we aimed to replicate our findings in validation cohorts. In samples from the treatment-naïve individuals in the IBSEN-III cohort the expansion of CD-associated meta-clonotypes was higher in individuals with CD relative to individuals with UC and SCs (**Fig. 2E**). The expansion of UC-associated meta-clonotypes was also higher in individuals with UC relative to SC but not to individuals with CD (**Fig. 2F**). Using another dataset from Rosati *et al.*^*14*^, CD-associated meta-clonotypes were expanded in individuals with CD relative to healthy controls, individuals with UC or with CRC (**Fig. 2G**). While UC-associated meta-clonotypes were significantly expanded in individuals with UC relative to healthy controls and CRC, but not individuals with CD (**Fig. 2H**).

### The expansion of CD-associated meta-clonotypes strongly correlates with disease location, behavior, and severity

Across the SPARC IBD cohort, the expansion of UC- and CD-associated meta-clonotypes correlated poorly with age (**Fig. S2A**, **S2B**) but moderately with years since diagnosis (**Fig. S2C**, **S2D)**. Particularly, CD-associated clonotypes showed a stronger correlation with years since diagnosis (ρ=0.144) relative to UC (ρ=-0.042). Additionally, the expansion of CD- and UC-associated clonotypes was not associated with the body mass index (**Fig. S2E, 2F**) and weakly with the biological sex, with females having a higher expansion of CD- and UC-associated meta-clonotypes (P=0.048 for CD and P=0.034 for UC; **Fig. S2G, S2H**).

The expansion of CD-associated meta-clonotypes was higher in ileal and ileocolonic CD relative to colonic CD (**Fig. 3A**), nonetheless, it still was higher in colonic CD than UC (**Fig. 3A**). The expansion of UC-associated meta-clonotypes was higher in individuals with UC relative to individuals with any form of CD (**Fig. 3B**). We replicated these findings using the IBSEN-III cohort, observing the highest expansion of CD-associated meta-clonotypes in ileocolonic CD and the lowest expansion in colonic CD (**Fig. 3C**). Nonetheless, UC-associated meta-clonotypes had a similar level of expansion in individuals with UC or CD (**Fig. 3D**). These observations, indicate that UC-associated meta-clonotypes might not be specific to UC *per se* but to colonic inflammation. Disease behavior strongly correlated with the expansion of CD-associated meta-clonotypes, specifically, the presence of a stricturing (B2), penetrating (B3) or both, *i.e*. B2/B3 disease behavior, was associated with a higher expansion relative a non-stricturing, non-penetrating disease course (B1) (**Fig. 3E**). Furthermore, it was higher in individuals with perianal fistulae relative to individuals without (**Fig. 3F**).

**Figure 3:**
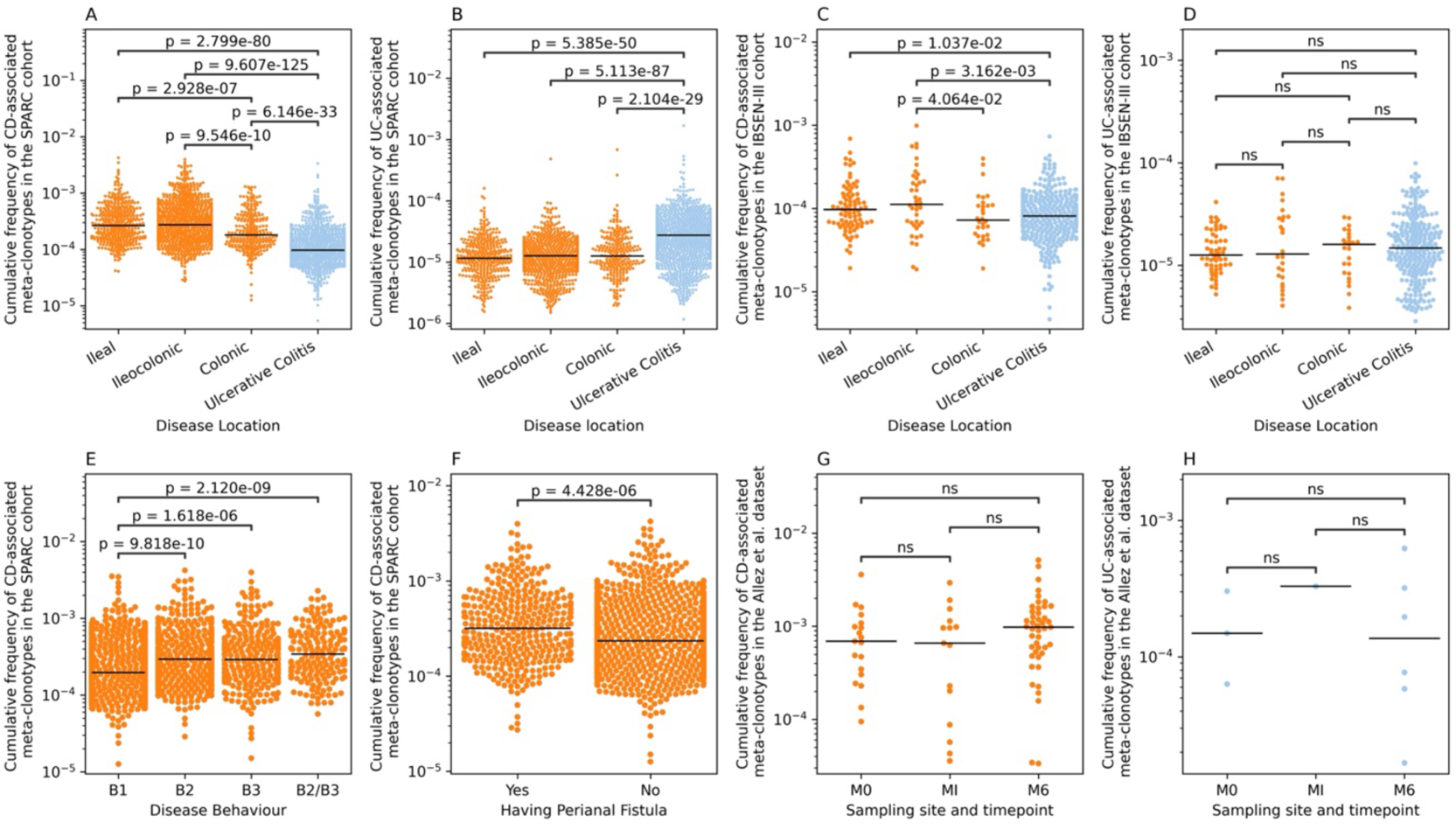
*The expansion of CD- and UC-associated meta-clonotypes in different clinical subtypes.* (**A**) shows the expansion of CD-associated meta-clonotypes in individuals from the SPARC IBD cohort with either ileal CD, ileocolonic CD, colonic CD, or ulcerative colitis, while (**B**) shows the expansion of UC-associated meta-clonotypes in the same phenotypic groups. (**C**) illustrates the expansion of CD-associated meta-clonotypes in treatment-naive individuals from the IBSEN-III cohort with ileal, ileocolonic, or colonic CD or UC, similarly, (**D**) shows the expansion of UC-associated meta-clonotypes in the same individuals. (**E**) depicts the expansion of CD-associated clonotypes in individuals with CD from the SPARC IBD cohort with different disease modifiers. (**F**) shows the expansion of CD-associated meta-clonotypes in individuals with and without perianal fistula. (**G**) shows the expansion of CD-associated meta-clonotypes in the inflamed ileum (M0) and proximal margin tissues (MI) of individuals with CD at the time of surgery, while (M6) represents the repertoire of biopsies collected from the same individuals 6 months post-operative. Similarly, (**H**) shows the expansion of UC-associated meta-clonotypes in the same phenotypic group as (**G**). The repertoire of samples used in (**G**) and (**H**) were obtained from Allez *et al.*^*25*^.

### CD- and UC-associated meta-clonotypes can be detected in the gut of individuals with IBD

Consequently, we aimed to investigate the expansion of these meta-clonotypes in the gut of individuals with IBD. Thus, we utilized the Rosati *et al.*^*14*^ dataset, which contains a bulk TRB repertoire of gut tissues from individuals with CD, UC, and CRC. CD-associated meta-clonotypes were significantly more expanded in the gut of CD relative to UC but at comparable levels to individuals with CRC (**Fig. S3A**). However, UC-associated meta-clonotypes showed comparable levels of expression across the three phenotypes (**Fig. S3B**), potentially because of the small sample size and the low expansion of UC-associated meta-clonotypes in general. Nonetheless, these findings indicate that a subset of the CD- and UC-associated clonotypes detected from blood are also observed in gut tissues.

Additionally, we used a previously published dataset by Allez *et al.*^*25*^ containing the TRB repertoire of inflamed (M0) and adjacent non-inflamed (MI) ileal tissues of individuals with ileal or ileocolonic CD undergoing ileal resection. The dataset^*25*^ also contained the repertoire of the neo-ileum 6 months post-operative (M6). While there was not any significant difference in the expansion of CD-associated meta-clonotypes in inflamed and non-inflamed tissues, CD-associated meta-clonotypes were detected in the local gut repertoire (**Fig. 3G**). Interestingly, the expansion of these clonotypes was not significantly different before and after surgery, suggesting that disease-associated clonotypes cannot be depleted or removed from the repertoire of individuals with IBD by surgery (**Fig. 3G**). UC-associated meta-clonotypes were not detected in the ileal immune repertoire of these individuals either before or after surgery (**Fig. 3H**), suggesting that UC-associated clonotypes have a lower presence in the ileum compared to the colon.

### CD- and UC- associated meta-clonotypes are mainly restricted to disease-associated HLA alleles and haplotypes

Given that conventional T cells are restricted to HLA proteins, we aimed to discover the cognate HLA proteins to which CD- and UC-associated meta-clonotypes preferentially bind. Thus, we utilized our recently published HLA-TCR dataset^*26*^, which contains ∼10^*6*^ pairs of TRB clonotypes and their associated HLA alleles. Out of the 908 UC-associated clonotypes, the cognate HLA partners for 67 TRB clonotypes could be resolved (**Fig. S4A**) with most of them being associated with the HLA-DRB1*15:01 allele (n= 21; ∼31.34%), corroborating our previous genetic fine mapping studies^*5*^. Multiple UC-associated clonotypes were restricted to other HLA alleles located on the same haplotype as HLA-DRB1*15:01, for example, HLA-B*07:02 (n=9), HLA-C*07:02 (n=11) and HLA-DQA1*01:02-DQB1*06:02 (n=14). Given the statistical nature by which these clonotypes are identified^*26*^, multiple clonotypes can be associated with different HLA alleles located on the same haplotype. Out of the 67 clonotypes with a cognate HLA allele, only 11 clonotypes were restricted to multiple HLA alleles, and the remaining clonotypes (n=56) were restricted to only one HLA allele.

Using the same analytical framework, we resolved the HLA restriction of 656 TRB clonotypes out of the 2,827 CD-associated clonotypes (**Fig. S4B**) with the majority being restricted to the HLA-DRB1*07:01 alleles (n=216; ∼33%). This allele has been genetically implicated in ileal CD^*11*^ and as a risk allele for CD in general in the HLA fine-mapping study by Goyette *et al.* (OR=1.14 for CD and OR=0.73 for UC)^*6*^. CD-associated clonotypes were also restricted to other HLA alleles that are located on the same haplotype as HLA-DRB1*07:01, for example, HLA-DQA1*02:01-DQB1*02:02 (n=150) and HLA-DQA1*02:01-DQB1*03:03 (n=131), potentially because of the strong linkage-disequilibrium observed among these HLA alleles. 438 clonotypes (66.76%) out of the 656 clonotypes were restricted to one HLA allele, *i.e.,* we observed 438 unique TRB-HLA pairs. Other clonotypes (n=218) were simultaneously restricted to multiple HLA alleles (mainly DRB1-DQA1-DQB1 haplotypes but in some cases extended C-B-DRB1-DQA1-DQB1-DPA1-DPB1 haplotypes).

To link TCR sequence similarity with HLA restriction, we performed a graph analysis on the identified disease-associated clonotypes (**Material and Methods**). Starting with UC-associated clonotypes, we observed multiple distinct clusters (**Fig. S5A**). Some of the clusters contain clonotypes that are restricted to disease-associated alleles such as HLA-DRB1*15:01 (**Fig. S5B**-**5C**), HLA-DQA1*01:03 (**Fig. S5D**) and HLA-DRB1*13:01^*27*^ (**Fig. S5D**). Two large-distinct clusters were not associated with any HLA alleles present in the database. This suggests that these clonotypes are either derived from unconventional T cells or are rarer in individuals without UC, and hence, we were not able to discover their cognate HLA alleles using our statistical analysis approach^*26*^.

The CD-associated meta-clonotypes showed a more complicated clustering pattern (**Fig. 4A**). The biggest cluster contained clonotypes restricted to the HLA-DRB1*07:01 allele and other HLA alleles located on the same haplotype (**Fig. 4B**). Multiple clusters were associated with other HLA-II proteins such as HLA-DPA1*03:01-DPB1*04:01 (**Fig. 4C**, **4E**), HLA-DQA1*01:02-DQB1*06:02 (**Fig. 4D**). Further, multiple distinct clusters were associated to non-DRB1*07:01 DRB1 proteins such as DRB1*11:01 (**Fig.4F, 4G**). This distinct clustering pattern, coupled with the restriction to known risk alleles, suggests that CD-associated clonotypes are responding to multiple antigens presented by these HLA alleles.

**Figure 4:**
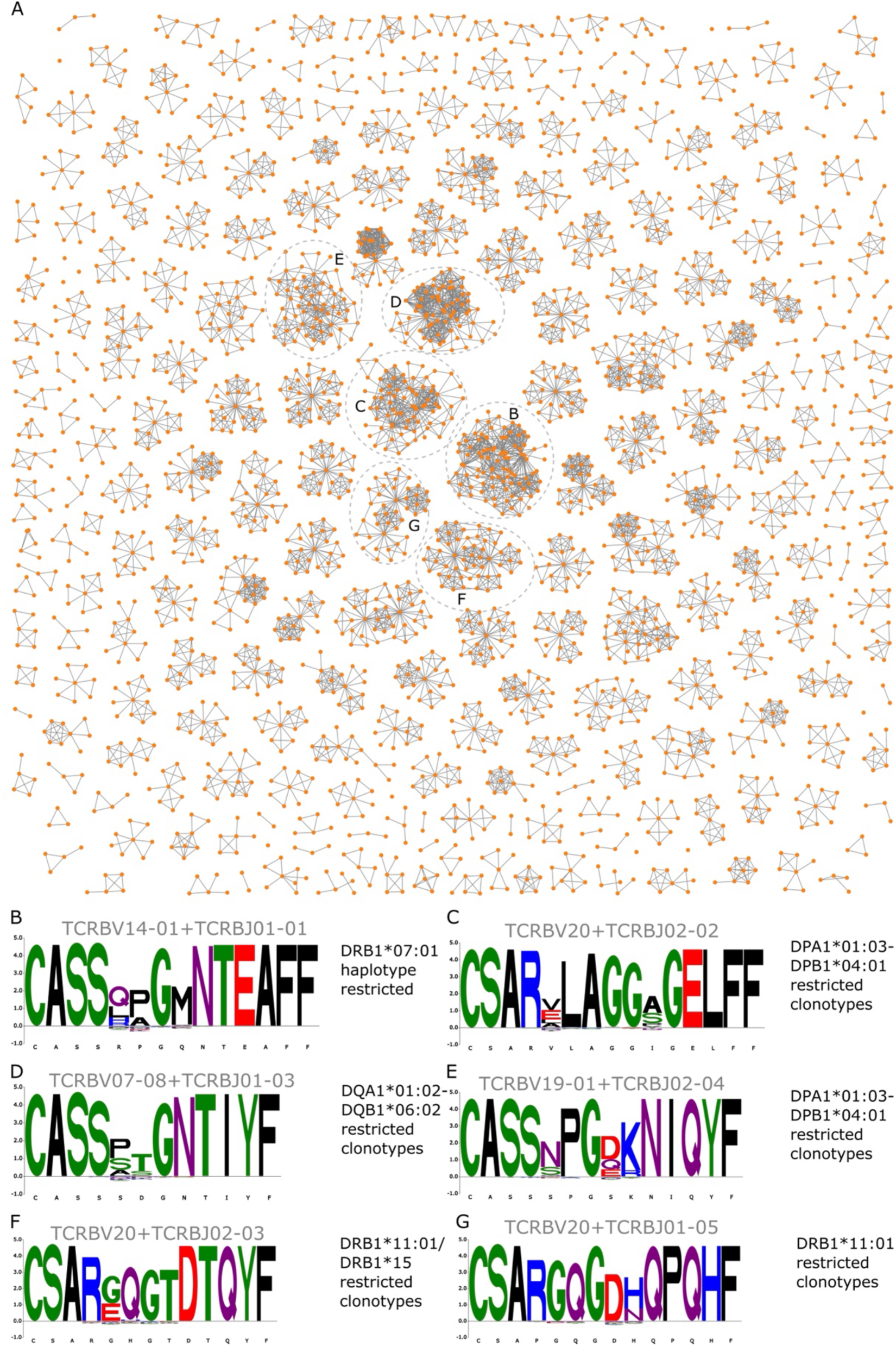
*Public immune responses associated with CD.* (**A**) Graph representation of CD-associated meta-clonotypes. Each node in the graph represents a clonotype, and edges represent similarities between these nodes. Specifically, two nodes are connected by an edge if they share the same V and J genes and there is a one-hamming distance between their CDR3 amino acid sequences. (**B**) shows the sequence motif of the biggest cluster containing multiple clonotypes that are restricted to HLA-DRB1*07:01 and its haplotype. (**C**) and (**E**) depict two distinct sequence motifs derived from two clusters that are restricted to the HLA-DPA1*01:03-DPB1*04:01 allele. (**D**) depicts the sequence motif of a cluster that harbors multiple clonotypes that are associated with HLA-DQA1*01:02-DQB1*06:02 clonotypes. Lastly, (**F**) and (**G**) depict the sequence motifs of two clusters that are associated with multiple HLA-DRB1 alleles, predominantly HLA-DRB1*11:01.

To identify potential antigens recognized by the disease-associated clonotypes, we queried public TCR-antigen databases, namely, VDJdb^*28*^ and McPAS^*29*^. Out of the 908 UC-associated clonotypes, only one clonotype was annotated in the database, targeting the self-protein, melanoma antigen recognized by T cells 1 (MLANA). For the 2,827 CD-associated clonotypes, only seven were annotated and were recognizing multiple viral epitopes, *e.g.,* EBV and CMV epitopes, and common intracellular bacteria such as *M. tuberculosis.* By overlapping these clonotypes with previously published fungi-specific clonotypes^*13*^, we annotated the antigenic specificities of 16 CD-associated clonotypes (**Fig. S6**) that predominately targeted *S. cerevisiae*, corroborating previous findings^*13*^. Nonetheless, the antigenic targets remain to be identified for most UC- and CD-associated meta-clonotypes.

### Identification of TRB-clonotypes associated with different CD and UC phenotypic subsets

Motivated by these findings, we were interested in identifying clonotypes that are associated with different CD and/or UC subphenotypes, *e.g.,* ileal vs colonic CD. Thus, we used the same statistical framework described above to identify meta-clonotypes associated with different clinical subphenotypes. This enabled us to identify 49 clonotypes arranged into 10 meta-clonotypes that were associated with ileal vs. colonic CD and ten clonotypes derived from two meta-clonotypes that were associated with ileocolonic vs. colonic CD. Furthermore, we identified 130 meta-clonotypes containing 1,318 clonotypes that were associated with ileal vs. ileocolonic CD. Lastly, 1,799 clonotypes derived from 216 meta-clonotypes were associated with colonic CD vs. UC.

The cumulative expansion of the 10 meta-clonotypes associated with ileal CD relative to colonic CD was significantly higher in individuals with ileal CD clonotypes relative to individuals with colonic CD (**Fig. 5A**). Although having identified only two meta-clonotypes that were higher in ileocolonic relative to colonic CD, the expansion of these meta-clonotypes was higher in individuals with ileocolonic CD (**Fig. 3B**). Similarly, the expansion of the 130 meta-clonotypes associated with ileal CD relative to colonic CD was higher in individuals with ileal CD relative to individual with ileocolonic CD (**Fig. 3C**). Lastly, the expansion of colonic CD-associated meta-clonotypes was higher in individuals with CD relative to individuals with UC (**Fig. 3D**). Our results support clinical observations that ileal CD is a distinct subtype of CD compared to colonic and ileocolonic CD, for example, ileal CD is associated with delayed diagnosis and increased risk for extraintestinal manifestations^*30*^.

**Figure 5:**
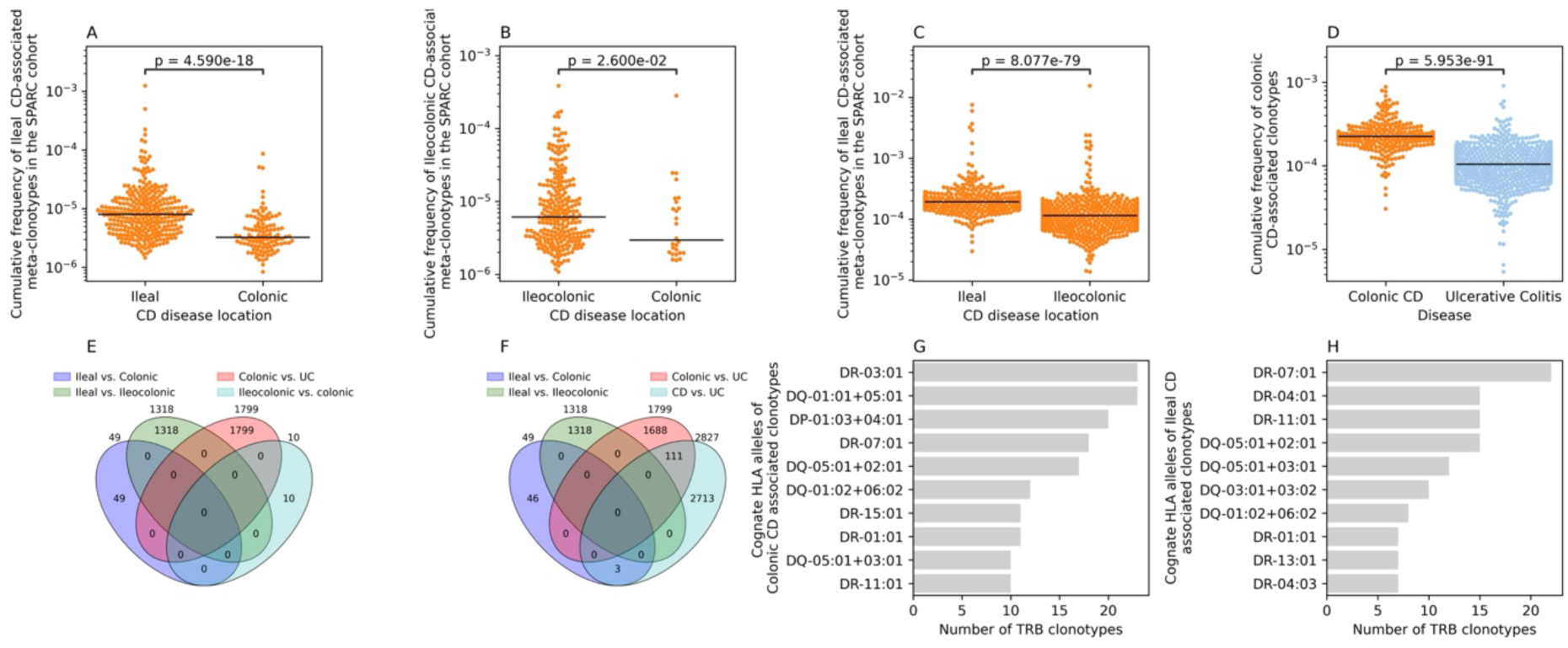
*Identification of TRB clonotypes that are associated with different CD subphenotypes.* These sets of subphenotype-associated clonotypes were discovered using a co-occurrence analysis that did not consider their expansion. Thus, to validate their expansion into their respective phenotypes, we compared their cumulative expansion among the different subphenotypes. (**A**) shows the expansion of ileal CD-associated meta-clonotypes (n=10) in individuals with ileal and colonic CD, while (**B**) shows the expansion of ileocolonic-associated meta-clonotypes in the TRB repertoire of individuals with ileocolonic and colonic CD. (**C**) illustrates the expansion of ileal CD associated meta-clonotypes identified by comparing the repertoire of individuals with ileal CD to individuals with ileocolonic CD. (**D**) shows the expansion of colonic CD-associated meta-clonotypes in the TRB repertoire of individuals with colonic CD or UC. (**E**) and (**F**) show the overlap in the identified clonotypes that are associated with different forms of IBD. (**G**) shows the HLA restriction of colonic CD-associated meta-clonotypes, while (**H**) illustrates the HLA restriction of ileal CD-associated meta-clonotypes.

The identified clonotypes were specific to each subphenotype and did not overlap among the different subtypes (**Fig. 3E**). Compared to the set of CD-associated clonotypes (n=327), only clonotypes that were associated with colonic-CD vs. UC showed an overlap (**Fig. 3F**). By analyzing the cognate HLA alleles^*26*^ of the colonic- and ileal-associated clonotypes we observed distinct HLA associations, with most colonic-CD specific clonotypes being associated with HLA-DRB1*03:01 (**Fig. 3G**). Most ileal CD-associated meta-clonotypes were restricted to HLA-DRB1*07:01 and HLA-DRB1*04:01 proteins (**Fig. 3H**). Thus, by analyzing the immune signature of different disease forms, we identified shared and subphenotype-specific antigenic exposures that elicit distinct immune responses in these different forms of IBD. TCR analyses may, therefore, be a helpful tool to define the different subtypes of IBD to better understand disease heterogeneity and to identify the relevant antigens in different subsets of the disease.

### Treatment and surgery have a minor impact on the expansion of CD- and UC-associated meta-clonotypes

Different therapeutic options for controlling IBD exist, ranging from conventional small-molecule-based therapies to biological-based therapies such as anti-TNF, *e.g.,* infliximab. A large proportion of the individuals in the SPARC IBD cohort were treated with anti-TNF, mainly infliximab and adalimumab, in addition to conventional therapy such as mesalamine and corticosteroid (**Table S3**). Surgeries are also commonly conducted in individuals with IBD to control the disease and to induce remission through partial or complete colectomy, ileal pouch-anal anastomosis, *i.e.,* J-pouch, or ileostomy. In the SPARC IBD cohort, about one-third of CD patients had at least one surgery, while about 10% of UC patients underwent surgery (**Table S4**). For CD patients, the most common surgeries were ileal and cecum resection, while it was complete colectomy for UC patients (**Table S4**).

To disentangle the effect of treatment from the underlying disease, we utilized the IBSEN-III cohort and compared the expansion of UC- and CD-associated meta-clonotypes in individuals with IBD before, *i.e.,* treatment-naive, and after treatment. Treatment trajectories did not influence the expansion of CD-associated meta-clonotypes in individuals with CD (**Fig. 6A**) and were associated with a slight reduction in the expansion of UC-associated meta-clonotypes in individuals with UC (**Fig. 6B**). Furthermore, neither the expansion of CD- nor UC-associated meta-clonotypes correlated with disease severity scoring (**Fig. 6C** & **6D**). Given the prospective nature of the SPARC IBD cohort, clinical data and immune profiles might not be temporally matched. Nonetheless, these observations indicate that the expansion of disease-associated clonotypes persisted under the different treatment trajectories.

**Figure 6:**
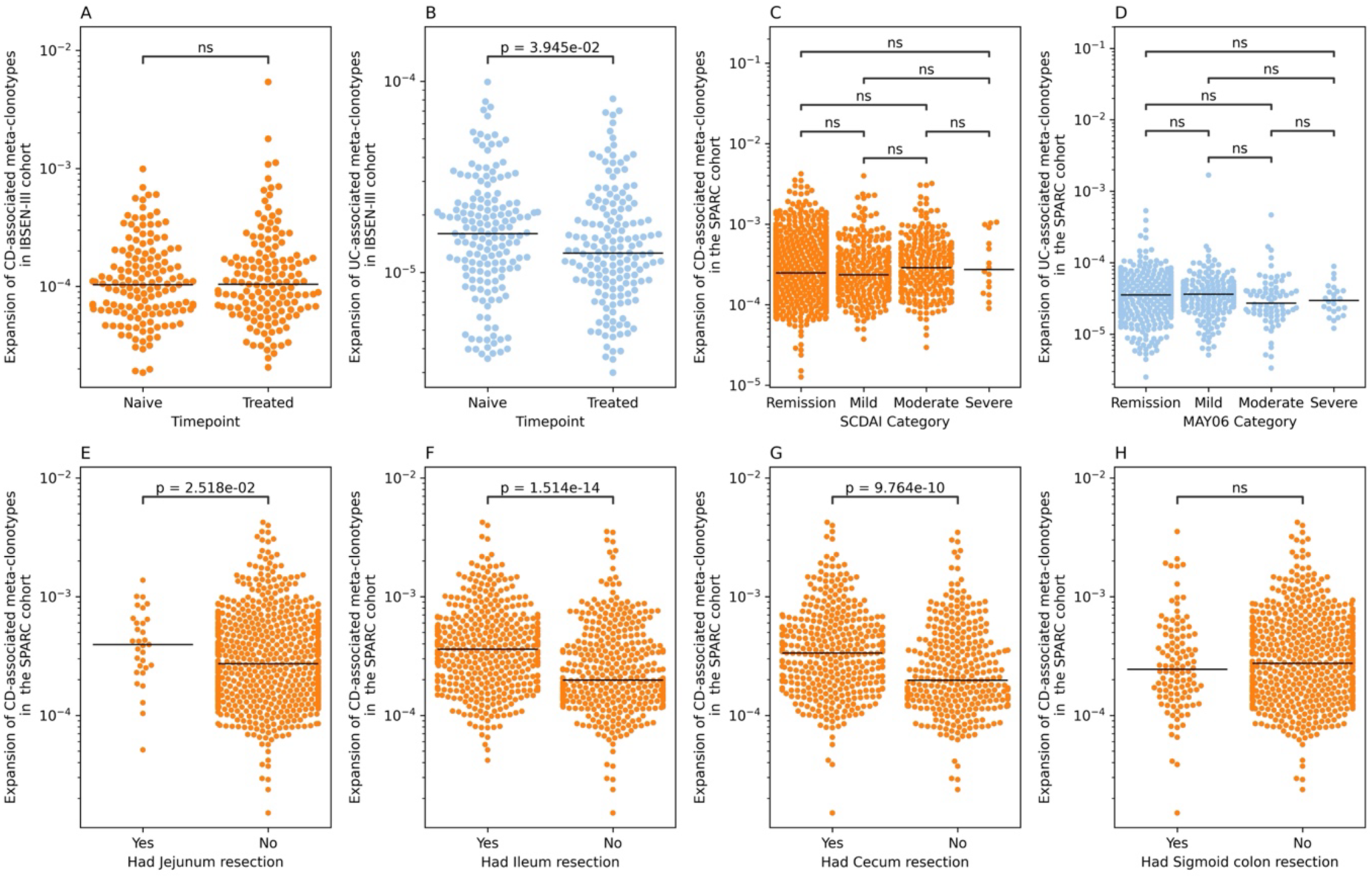
*The impact of medication and surgery on the expansion of CD and UC-associated meta-clonotypes*. (**A**) shows the expansion of CD-associated meta-clonotypes in the same individuals with CD before any medication and at the time of diagnosis (“Naive”) and one-year after treatment (“Treated”). The same relationship is depicted in (**B**) but for UC-associated meta-clonotypes in individuals with UC. In (**A**) and (**B**), the Wilcoxon test was used to compare the expansion of CD- and UC-associated meta-clonotypes in treatment-naive and treated individuals. (**C**) depicts the expansion of CD-associated meta-clonotypes in individuals with CD against their different disease activity statuses as defined by the SCDAI score, while (**D**) shows the expansion of UC-associated meta-clonotypes in individuals with UC against their disease activity statuses as defined by the Mayo 6 score. (**E**) illustrates the expansion of CD-associated meta-clonotypes in individuals with CD that either underwent jejunum resection or not, while (**F**) shows the same relationship in individuals with ileum resection. (**G**) and (**H**) show the expansion of CD- associated meta-clonotypes in individuals with CD who underwent different colon resection surgeries, specifically, cecum resection (**G**) and sigmoid resection (**H**).

The expansion of CD-associated meta-clonotypes was significantly higher in patients who underwent either jejunal (**Fig. 6B**) or ileal resection (**Fig. 6F**). While this might be confounded by the disease subtype, *e.g.,* ileal CD where the expansion is higher or other complications such as stricturing or penetrating behavior, the persistent high expansion of these clonotypes indicates that the immune signature of the disease persisted even after removing parts of the affected organ. Also, individuals who underwent cecum resection had a higher expansion of CD-associated meta-clonotypes relative to individuals who did not undergo cecum resection (**Fig. 6G**). Nonetheless, individuals with and without sigmoid colon resection showed comparable levels of CD-associated meta-clonotypes expansion (**Fig. 6H**). The expansion of UC-associated meta-clonotypes was comparable in individuals with UC who either underwent or did not go cecum, sigmoid, or rectal resection, as well as complete colectomy (**Fig. S7**). These analyses suggest a strong interaction between disease location and a persisting, continuous immune response, particularly in CD. Specifically, inflammation involving the ileum and cecum seems to induce a robust expansion of different clonotypes that persist after the partial removal of the inflamed tissue.

## Discussion

The T cell repertoire contains T memory cells that encode previous and ongoing antigenic exposures. While the immune repertoire of individuals with IBD has been previously studied^*25, 31–33*^, these studies have focused on a smaller number of individuals (n<100). By comparing the immune repertoire of 1,890 individuals with CD to that of 914 individuals with UC, we were able to identify clonotypes that were associated with either form of IBD. The expansion of CD-associated clonotypes was significantly higher in individuals with CD relative to individuals with UC and healthy controls across the two validation datasets. This indicated that the identified set of CD-associated meta-clonotypes is a generalizable set of clonotypes that might be implicated in CD pathogenesis. On the other hand, UC-associated meta-clonotypes showed lower levels of expansion in the blood relative to CD-associated meta-clonotypes. Using the two independent validation datasets, we observed that the expansion of UC-associated meta-clonotypes is higher in individuals with UC and CD relative to healthy controls and CRC, suggesting that these clonotypes might be markers of colonic inflammation and not specifically UC. However, the replication of our findings in cohorts from Germany and Norway, which were profiled using different TCR-repertoire profiling technology from RNA instead of DNA, highlights the reproducibility and the strong involvement of these clonotypes in the pathogenesis of IBD. Furthermore, these clonotypes are not expanded as a direct or an indirect result of therapeutic interventions because of their significant expansion in treatment-naive individuals from the IBSEN-III cohort.

Most of the herein identified clonotypes were specific to HLA-II alleles, particularly several HLA-DRB1 alleles, with most CD-associated meta-clonotypes being restricted to HLA-DRB1*07:01 and UC-associated meta-clonotypes to HLA-DRB1*15:01. We were not able to decode the antigenic specificities of these clonotypes employing data from public databases^*28, 29*^ which contain mainly HLA-I presented antigens. However, by resolving the antigenic specificities of these clonotypes in the future, a better understanding of antigens driving or causing the disease can be obtained. Different experimental techniques could be utilized to identify the antigenic targets of CD- and UC-associated meta-clonotypes, such as yeast-^*34, 35*^ and phage-display^*36*^ based methods.

A subset of CD-associated clonotypes were targeting fungal antigens, predominantly *S. cerevisiae,* corroborating previous findings implicating fungal antigens in driving CD^*13*^. Nonetheless, anti-fungal responses represent a smaller subset of CD-associated meta-clonotypes (16 out of 2,827 CD-associated clonotypes), suggesting that besides fungal antigens, multiple yet-to-be-identified antigens are driving the disease. Although we were able to resolve the cognate HLA alleles of these clonotypes statistically^*26*^, we were not able to do this for a subset of disease-associated clonotypes. This subset can be rarer in non-IBD patients, and consequently, we were not able to identify their cognate HLA alleles using our statistical approach, which is based on a reference dataset that also contains healthy controls and individuals with PSC^*26*^. Another possibility is that these clonotypes can be assigned to unconventional T cells, such as CAIT cells^*14*^ or other yet-to-be-characterized populations of unconventional T cells that may play a role in IBD pathogenesis.

The larger number of CD-associated meta-clonotypes and their higher expansion in the blood relative to UC-associated clonotypes corroborates previous findings on the stronger immune signature of CD relative to UC^*37*^. Within the different subphenotypes of CD, ileal and ileocolonic CD had a higher expansion relative to colonic CD. These observations suggest an interaction between the location of the inflammation and the generated immune response, where inflammation at the end of the small intestine and the beginning of the colon appears to generate a stronger immune response. This can be attributed to differences in the composition of immune cells across the small intestine and colon^*38*^; for example, most Peyer’s patches are in the terminal ileum^*39*^.

A key finding of our study was the persistent expansion of CD-associated meta-clonotypes in individuals with CD who underwent surgery, particularly of the jejunum, ileum, and cecum, indicating that CD-associated clonotypes persist even after the removal of affected tissues, potentially explaining disease recurrence and flares that happen in patients after surgery. We validated this finding using the dataset of Allez *et al.*^*25*^, showing that the expansion of CD-associated clonotypes in ileal tissues is comparable before and after ileal resection surgeries. This suggests that either disease-associated clonotypes are primed in a distinct anatomical location other than the affected tissues or that these priming sites act as a reservoir of disease-associated clonotypes. Alternatively, these findings might indicate that there is a persistent antigenic exposure that stimulates the formation of these T-cell responses. Disentangling the cause of this apparent persistent expansion of CD-associated clonotypes may have a significant impact on developing novel therapies for IBD. For example, if it is mainly because of a persistent antigenic exposure, then identifying and removing this antigenic exposure may be the best therapeutic strategy. Through the targeted depletion of ankylosing spondylitis-associated clonotypes, Britanova and colleagues^*40*^ were able to develop antibodies that introduce remission and a reversal of ankylosing spondylitis symptoms. Alternatively, if these CD-associated clonotypes have been formed due to a previous antigenic exposure and are then maintained because of either cross-reactivity to a self-protein or the gut microbiome, then a targeted depletion of these clonotypes might represent the best therapeutic strategy.

In conclusion, by identifying the antigenic targets of the discovered CD- or UC-associated clonotypes, a better understanding of immune processes and antigens involved in IBD can be obtained. Furthermore, these clonotypes might be a promising target for novel therapeutic options to treat and control CD and UC.

## Material and Methods

### Study design

The SPARC IBD cohort contains the TRB repertoire (**Material and Methods**) of 1,890 and 914 individuals with CD and UC, respectively, together with detailed information regarding surgeries, patient-reported outcomes, and treatments (**Table S1**). Using the TRB repertoire of these 2,804 individuals, we used the statistical framework described by Emerson *et al*.^*19*^ to identify clonotypes that are statistically associated with CD or UC (**Material and Methods** & **Fig. 1A**). To validate the identified clonotypes, we profiled the TRB repertoire of samples from the “Inflammatory bowel disease in South-Eastern Norway III (IBSEN-III)” cohort^*41*^ (**Material and Methods; Fig. 1B**). This cohort contains samples derived from treatment-naive and treated individuals with CD, UC, or symptoms of IBD without supporting endoscopic or radiological findings, *i.e.,* symptomatic controls (SC) (**Table S2**). For a subset of individuals with CD or UC, the TRB repertoire was again profiled one-year post-diagnosis, *i.e.,* treated individuals, which enabled us to quantify the impact of therapeutic interventions on the expansion of CD- or UC-associated clonotypes (**Fig. 1B**). We further validated our findings (**Fig. 1C**) using a previously published dataset^*14*^ containing the TRB repertoire of 100 healthy controls and 118, 47 and 12 individuals with CD, UC or colorectal carcinoma (CRC), respectively, from Germany.

### Profiling the T cell repertoire of the SPARC IBD cohort

The SPARC IBD cohort is a US-based cohort of individuals with IBD that is maintained by the Crohn’s & Colitis Foundation IBD Plexus program for research purposes^*24*^. The TRB repertoire was profiled by the foundation using ImmuneSEQ (Adaptive biotechnologies), as discussed previously^*20, 42*^.

### Profiling the T cell repertoire of the IBSEN-III cohort

PAXgene tubes were collected at the time of diagnosis, *i.e.,* the treatment-naive status, and at a one-year clinical follow-up, *i.e.*, treated samples. To profile the T cell receptor, RNA was extracted from the PAXgene tubes using the QIAcube system (QIAGEN). Subsequently, 300ng of RNA were used to profile the blood repertoire using a commercially available kit from (MiLaboratories) according to the Manufacturer’s instructions. After preparing the sequencing libraries using MiLaboratories’s kits and Illumina Unique Dual index system, different samples were pooled together and sequenced on an S4 flow cell using a NovaSeq 6000 sequencer. For each sample, we aimed for ∼5 million reads (150bp X2). After sequencing and sample demultiplexing, we utilized MiXCR^*43*^ with the default settings to assemble clonotypes and quantify their expansion in each sample.

### Secondary processing of the identified clonotypes

After identifying clonotypes from sequencing reads either from the SPARC IBD or the IBSEN-III cohort, we processed the identified clonotypes by removing non-productive clonotypes, *i.e.,* clonotypes containing a stop-codon or a frameshift, as they do not encode for a functional TRB chain. Due to the degeneracy of the genetic code, different CDR3 nucleotide sequences will encode for the same amino acid, *i.e.,* different V(D)J recombination at the DNA level will encode for the same TRB chain at the protein level. Given that we focused on the expressed proteins that form the TCR, we grouped all V(D)J recombination products that utilize the same V and J genes and encode the same CDR3 amino acid sequences together and summed their expansions. Hence, in all the analyses, a clonotype is a productive, unique recombination composite of a unique V and J gene combination in conjunction with a unique CDR3 amino acid sequence.

### Identifying disease-associated clonotypes

To identify disease-associated clonotypes, we utilized the same framework described by Emerson *et al.*^*19*^, where we started by identifying public clonotypes, *i.e.,* clonotypes identified in two or more individuals. Subsequently, we compared the count of individuals with either CD or UC carrying these clonotypes to individuals without CD or UC who do not carry these clonotypes using the Fisher’s exact test (FET). We utilized two one-sided FETs to identify clonotypes that are associated with either CD or UC. After calculating the association P-value for each public clonotype, we defined disease-associated clonotypes as clonotypes with an association P-value < 1x10^-4^. The same paradigm was also utilized to identify clonotypes associated with different subsets of the disease, such as ileal, ileocolonic, or colonic CD.

### Extending disease-associated clonotypes using seeded clustering

To obtain a comprehensive set of disease-associated TRB clonotypes, we performed seeded clustering. Seeded clustering is based on three steps: first, seed identification; second, extended search; and lastly, set purification. In the first step, the framework proposed by Emerson *et al.*^*19*^ is utilized to identify clonotypes that are associated with the disease, as discussed above. In the extended search, for each seed, the union of all samples’ repertoires is queued to identify clonotypes that have the same V and J genes as the seed and have a Levenshtein distance of one from the seed. After performing the extended search for each seed, multiple highly similar clonotypes will be identified. We refer to these clonotypes as the seed-associated clonotype set. Lastly, purification is conducted to ensure that each member of the seed-associated clonotype set is implicated in the disease or associated with the trait under-investigation. During the purification step, we compared the strength of disease association, as measured by the FET, between the seed alone and the seed and each member of the seed-associated clonotype set. If adding a member to the leading seed results in a reduction of the disease association P-value, *i.e.,* an increase in the numerical value of FET’s P-value, then this member is removed from the set of seed-associated clonotypes; otherwise, it is kept. After purifying the set of seed-associated clonotypes, this set, and the seed are treated as one meta-clonotype that is associated with the disease.

### Network analysis and visualization

To perform network analysis, we treated each sequence as a node in a graph; an edge between two nodes is drawn if both nodes share the same V and J genes, and the distance between their CDR3 amino acid sequences is at most one-hamming distance. Visualization was conducted using Cystoscape^*44*^.

## Author contributions

H.E. and A.F. conceived and designed the study. A.K.H.M performed the data curation and processing of the SPARC IBD clinical data. A.K.H.M, E.E.K, and V.K. prepared the TRB NGS libraries from the IBSEN-III cohort. C.O., G.P., M.B.B., P.R., S.A., T.E.D., V.A.K., J.R.H., and M.L.H. collected the biomaterial from the IBSEN-III cohort, in addition to collecting, storing, and processing clinical and metadata. H.E. and A.K.H.M. made the figures. H.E. and A.K.H.M. performed the analyses and wrote the first draft of the manuscript with input from all co-authors. All authors read and approved the final version of the manuscript.

## Data availability

Two main datasets were utilized in the current study, the first is the TRB repertoire of the “A Study of a Prospective Adult Research Cohort with IBD (SPARC IBD)” cohort of the Crohn’s & Colitis Foundation IBD Plexus program. The second cohort is the TRB repertoire of the “Inflammatory bowel disease in South-Eastern Norway III (IBSEN-III)”. The US-based SPARC IBD dataset is available upon approved application to the Crohn’s & Colitis Foundation IBD Plexus Program (https://www.crohnscolitisfoundation.org/ibd-plexus). Regarding the IBSEN III dataset, institutional data privacy regulations prohibit the deposition of individual-level data to public repositories. Participant written consent also does not cover public sharing of data for use for unknown purposes. Upon contact with Marte Lie Høivik an institutional data transfer agreement can be established and data shared if the aims of data use are covered by ethical approval and patient consent. The procedure will involve an update to the ethical approval as well as review by legal departments at both institutions, and the process will typically take one to two months from initial contact. The analytical code and software used in the current study are based on publicly available tools and custom scripts developed in Python, as described in the Material and Methods section.

## Ethical approval

The IBSEN III study was approved by the South-Eastern Regional Committee for Medical and Health Research Ethics (Ref 2015/946-3) and performed in accordance with the Declaration of Helsinki. All patients signed an informed consent form prior inclusion in this study and the data were stored in services for sensitive data (TSD) at the University of Oslo.

## Acknowledgments

The project was funded by the EU Horizon Europe Program grant *miGut-Health: Personalized blueprint of intestinal health* (101095470). Additionally, the project received infrastructure support from the German Research Foundation (DFG) Research Unit 5042: miTarget – The Microbiome as a Therapeutic Target in Inflammatory Bowel Diseases and from the DFG Cluster of Excellence 2167 “Precision Medicine in Chronic Inflammation (PMI)”. A.K.H.M. is funded by the DFG Collaborative Research Unit 1526 „Pathomechanisms of Antibody-mediated Autoimmunity (PANTAU) – Insights from Pemphigoid Diseases. The funders had no role in study design, data collection and analysis, decision to publish, or preparation of the manuscript. We are grateful to the Crohn’s & Colitis Foundation IBD Plexus program for providing us with the T cell repertoire profiles of the SPARC IBD cohort. We would also like to thank Michael Wittig (Institute of Clinical Molecular Biology, Kiel University and University Hospital Schleswig-Holstein, Kiel, Germany) for helping with interpreting and processing the genotyping datasets. We are also thankful to Sören Franzenburg, Janina Fuß, Rebekka Kraemer, Nicole Braun, Maria Eloina Figuera Basso, Anja Tanck, Xiaoli Yi, Tanja Wesse, Yewgenia Dolshanskaya, Melanie Vollstedt and Cathrin John-Klaua from the Institute of Clinical Molecular Biology (Kiel University and University Hospital Schleswig-Holstein, Kiel, Germany) for their help with profiling the TRB repertoire of the IBSEN-III cohort. Figure 1 was Created in https://BioRender.com Lastly, we would like to thank the IBSEN-III study group for their efforts in collecting and providing the samples. IBSEN-III study group:

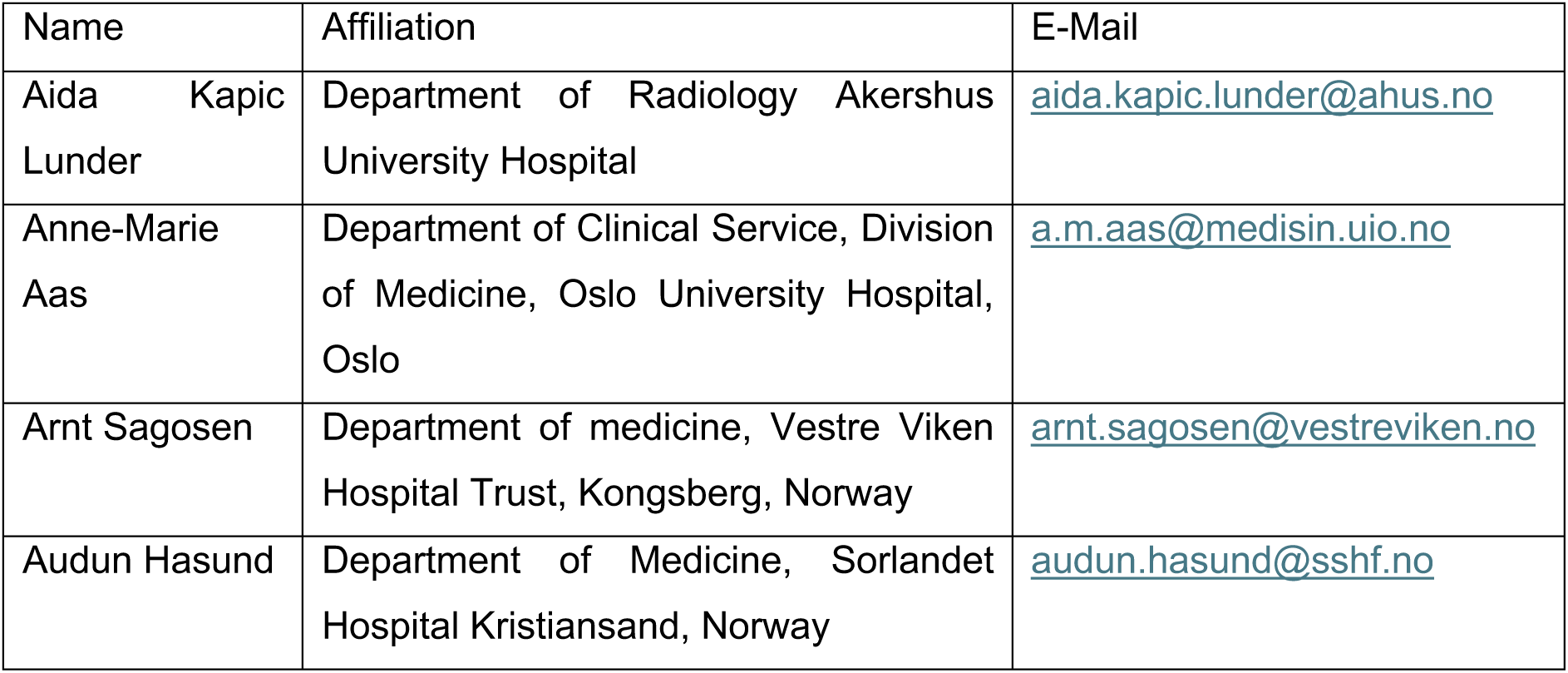
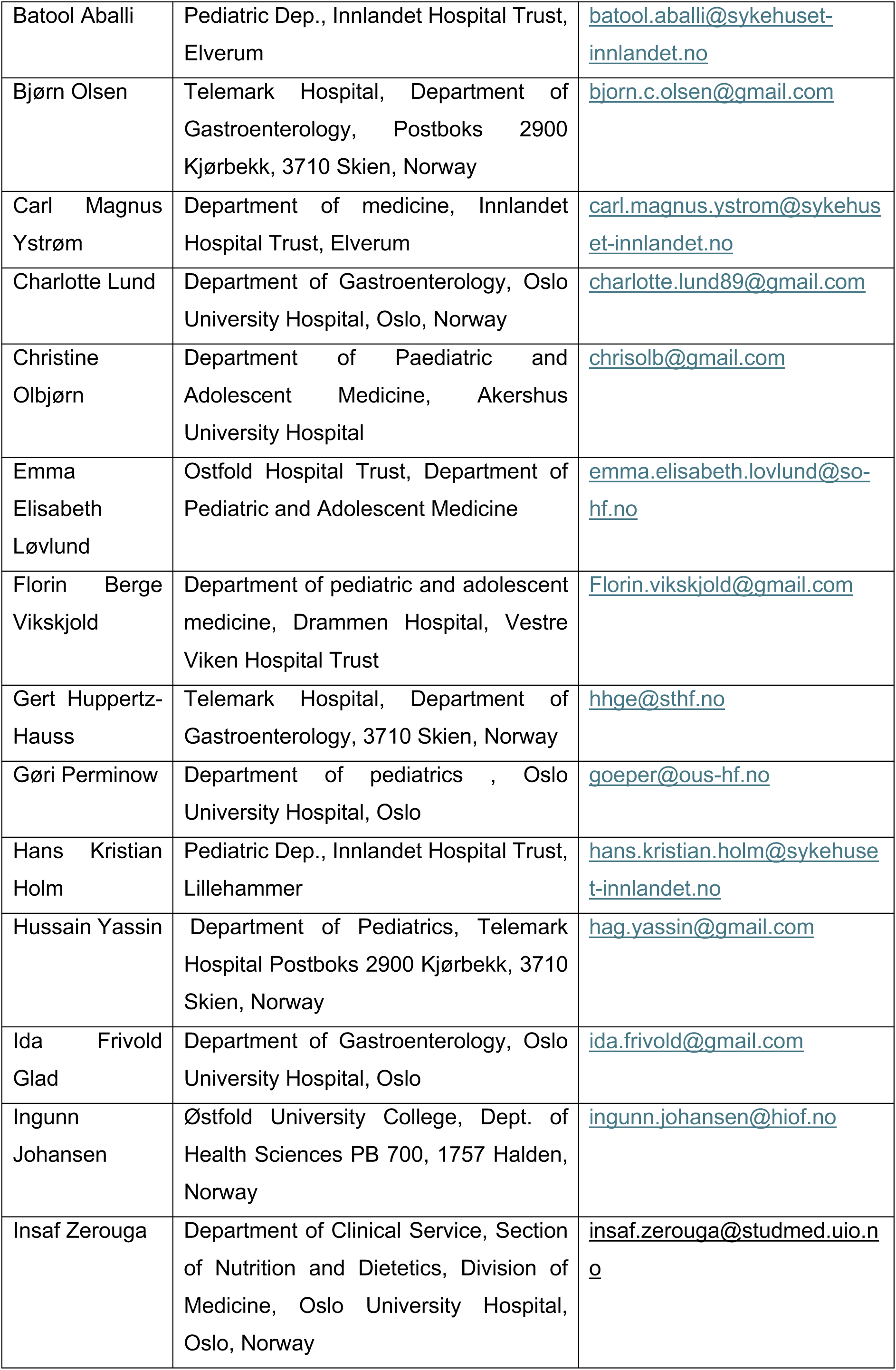
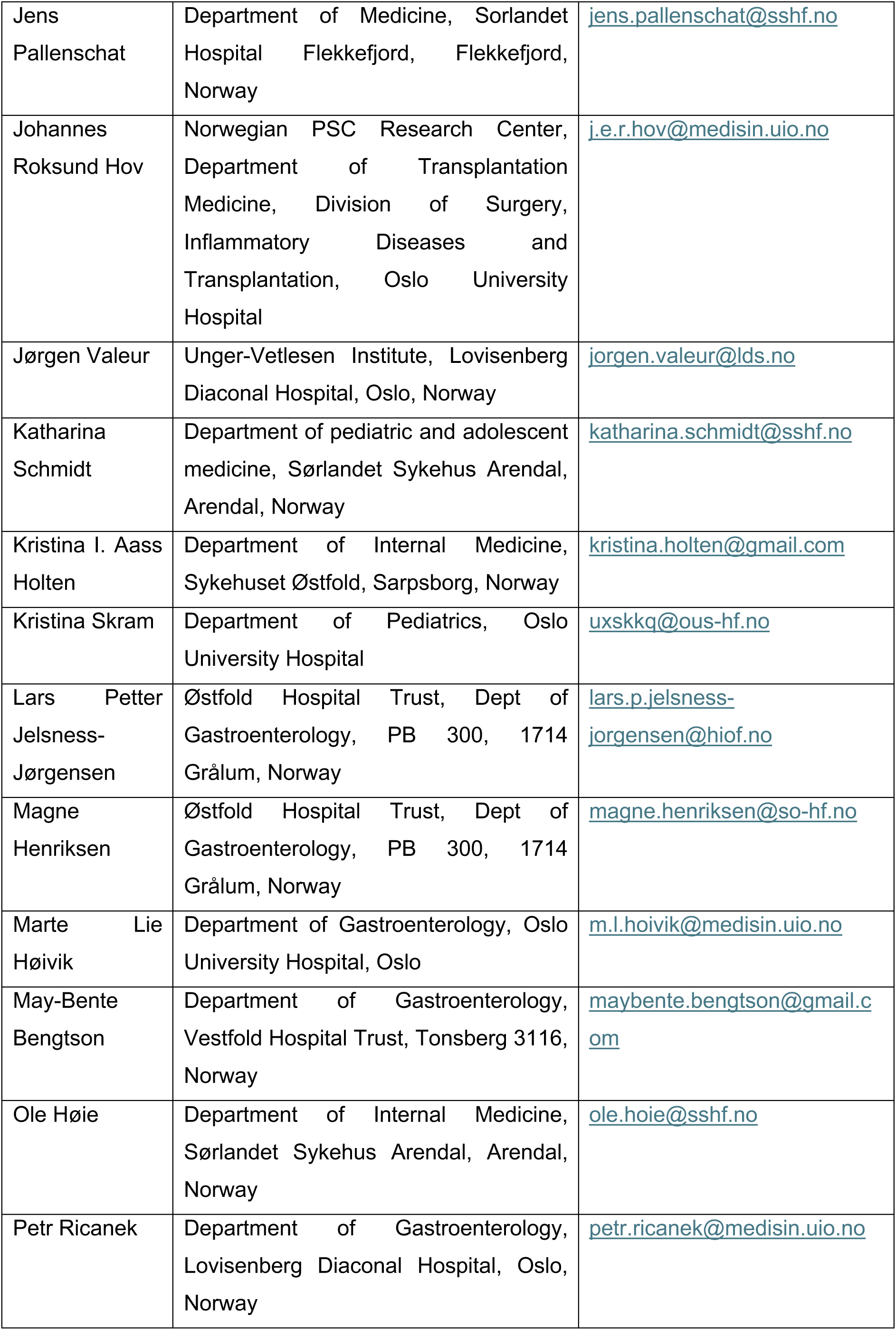
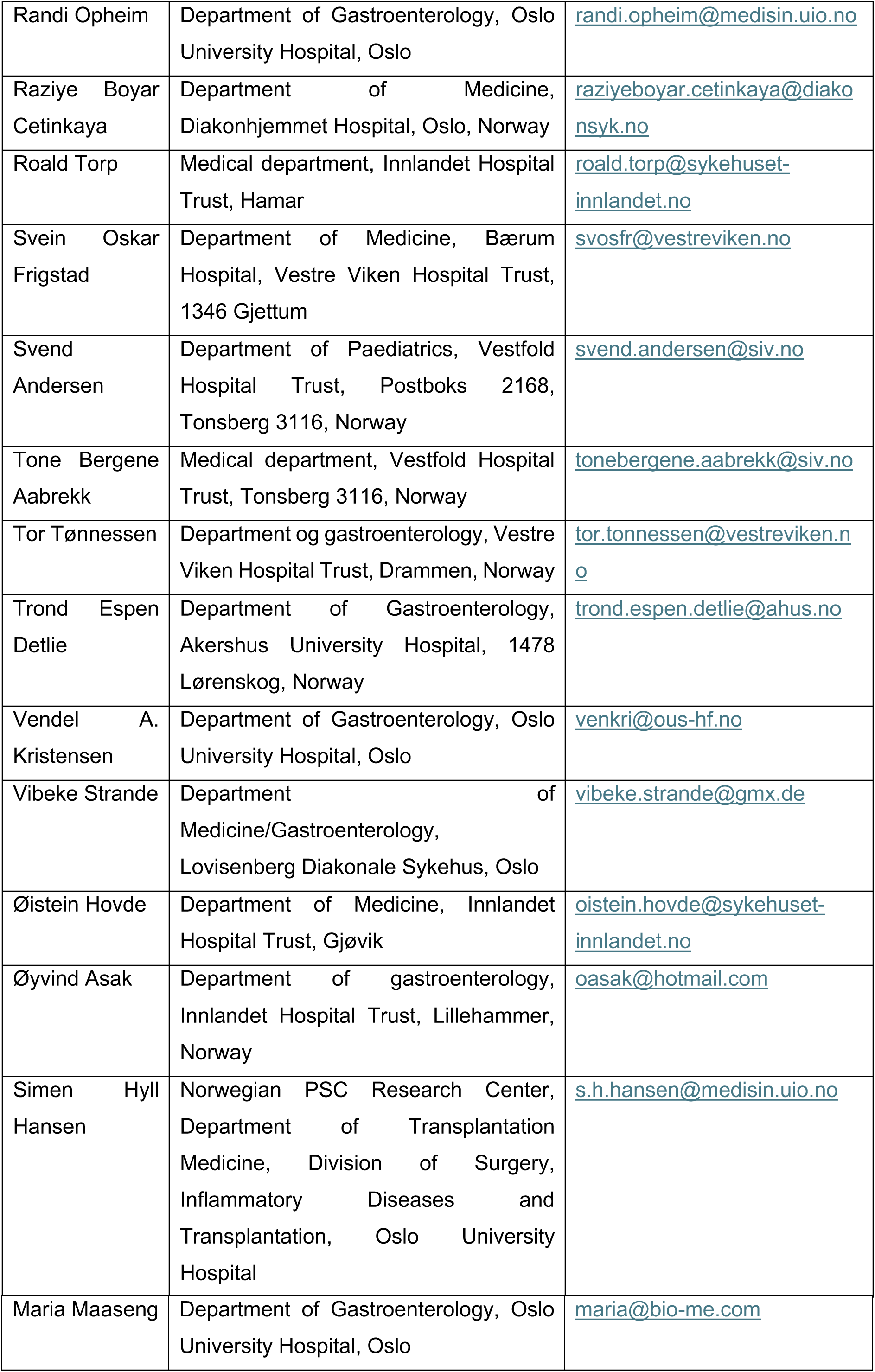
Descriptive comparisons of traditional septum thickness measurements by three independent observers.

## References

1. Yadav, P. et al. Genetic Factors Interact With Tobacco Smoke to Modify Risk for Inflammatory Bowel Disease in Humans and Mice. Gastroenterology 153, 550–565 (2017).

2. Shan, Y., Lee, M. & Chang, E. B. The gut microbiome and inflammatory bowel diseases. Annu Rev Med 73, 455–468 (2022).

3. Jostins, L. et al. Host–microbe interactions have shaped the genetic architecture of inflammatory bowel disease. Nature 491, 119–124 (2012).

4. Ebert, A. C. et al. Risk of inflammatory bowel disease following hospitalisation with infectious mononucleosis: nationwide cohort study from Denmark. Nat Commun 15, 8383 (2024).

5. Degenhardt, F. et al. Trans-ethnic analysis of the human leukocyte antigen region for ulcerative colitis reveals shared but also ethnicity-specific disease associations. Hum Mol Genet (2021) doi:10.1093/hmg/ddab017.

6. Goyette, P. et al. High-density mapping of the MHC identifies a shared role for HLA-DRB1*01:03 in inflammatory bowel diseases and heterozygous advantage in ulcerative colitis. Nat Genet 47, 172–179 (2015).

7. Franke, A. et al. Genome-wide meta-analysis increases to 71 the number of confirmed Crohn’s disease susceptibility loci. Nat Genet 42, 1118–1125 (2010).

8. Ogura, Y. et al. A frameshift mutation in NOD2 associated with susceptibility to Crohn’s disease. Nature 411, 603–606 (2001).

9. Hugot, J.-P. et al. Association of NOD2 leucine-rich repeat variants with susceptibility to Crohn’s disease. Nature 411, 599–603 (2001).

10. Hampe, J. et al. A genome-wide association scan of nonsynonymous SNPs identifies a susceptibility variant for Crohn disease in ATG16L1. Nat Genet 39, 207–211 (2007).

11. Ahmad, T. et al. The molecular classification of the clinical manifestations of Crohn’s disease. Gastroenterology 122, 854–866 (2002).

12. Uchida, A. M. et al. Escherichia coli–Specific CD4+ T Cells Have Public T-Cell Receptors and Low Interleukin 10 Production in Crohn’s Disease. Cell Mol Gastroenterol Hepatol 10, 507–526 (2020).

13. Martini, G. R. et al. Selection of cross-reactive T cells by commensal and food-derived yeasts drives cytotoxic TH1 cell responses in Crohn’s disease. Nat Med 29, 2602–2614 (2023).

14. Rosati, E. et al. A novel unconventional T cell population enriched in Crohn’s disease. Gut 71, 2194 LP – 2204 (2022).

15. Bashford-Rogers, R. J. M. et al. Analysis of the B cell receptor repertoire in six immune-mediated diseases. Nature 574, 122–126 (2019).

16. Lodes, M. J. et al. Bacterial flagellin is a dominant antigen in Crohn disease. J Clin Invest 113, 1296–1306 (2004).

17. Bourgonje, A. R., Hörstke, N. V, Fehringer, M., Innocenti, G. & Vogl, T. Systemic antibody responses against gut microbiota flagellins implicate shared and divergent immune reactivity in Crohn’s disease and chronic fatigue syndrome. Microbiome 12, 141 (2024).

18. Mahdy, A. K. H. et al. Bulk T cell repertoire sequencing (TCR-Seq) is a powerful technology for understanding inflammation-mediated diseases. J Autoimmun 149, 103337 (2024).

19. Emerson, R. O. et al. Immunosequencing identifies signatures of cytomegalovirus exposure history and HLA-mediated effects on the T cell repertoire. Nat Genet 49, 659–665 (2017).

20. Gittelman, R. M., et al. Longitudinal analysis of T cell receptor repertoires reveals shared patterns of antigen-specific response to SARS-CoV-2 infection. JCI Insight 7, (2022).

21. Greissl, J. et al. Immunosequencing of the T-Cell Receptor Repertoire Reveals Signatures Specific for Identification and Characterization of Early Lyme Disease. medRxiv 2021.07.30.21261353 (2022) doi:10.1101/2021.07.30.21261353.

22. Rawat, P. et al. Identification of a type 1 diabetes-associated T cell receptor repertoire signature from the human peripheral blood. medRxiv 2024.12.10.24318751 (2024) doi:10.1101/2024.12.10.24318751.

23. Pesesky, M. et al. Antigen-driven expansion of public clonal T cell populations in inflammatory bowel diseases. J Crohns Colitis jjaf048 (2025) doi:10.1093/ecco-jcc/jjaf048.

24. Raffals, L. E. et al. The Development and Initial Findings of A Study of a Prospective Adult Research Cohort with Inflammatory Bowel Disease (SPARC IBD). Inflamm Bowel Dis 28, 192–199 (2022).

25. Allez, M. et al. T cell clonal expansions in ileal Crohn’s disease are associated with smoking behaviour and postoperative recurrence. Gut 68, 1961–1970 (2019).

26. ElAbd, H. et al. Decoding the restriction of T cell receptors to human leukocyte antigen alleles using statistical learning. bioRxiv 2022–2025 (2025).

27. Cleynen, I. et al. Inherited determinants of Crohn’s disease and ulcerative colitis phenotypes: a genetic association study. The Lancet 387, 156–167 (2016).

28. Goncharov, M. et al. VDJdb in the pandemic era: a compendium of T cell receptors specific for SARS-CoV-2. Nat Methods 19, 1017–1019 (2022).

29. Tickotsky, N., Sagiv, T., Prilusky, J., Shifrut, E. & Friedman, N. McPAS-TCR: a manually curated catalogue of pathology-associated T cell receptor sequences. Bioinformatics 33, 2924–2929 (2017).

30. Dulai, P. S. et al. Should We Divide Crohn&#x2019;s Disease Into Ileum-Dominant and Isolated Colonic Diseases? Clinical Gastroenterology and Hepatology 17, 2634–2643 (2019).

31. Chapman, C. G. et al. Characterization of T-cell Receptor Repertoire in Inflamed Tissues of Patients with Crohn’s Disease Through Deep Sequencing. Inflamm Bowel Dis 22, 1275–1285 (2016).

32. Saravanarajan, K. et al. Genomic profiling of intestinal T-cell receptor repertoires in inflammatory bowel disease. Genes Immun 21, 109–118 (2020).

33. Werner, L. et al. Altered T cell receptor beta repertoire patterns in pediatric ulcerative colitis. Clin Exp Immunol 196, 1–11 (2019).

34. Wang, L. & Lan, X. Rapid screening of TCR-pMHC interactions by the YAMTAD system. Cell Discov 8, 30 (2022).

35. Birnbaum, M. E. et al. Deconstructing the Peptide-MHC Specificity of T Cell Recognition. Cell 157, 1073–1087 (2014).

36. Ch’ng, A. C. W., Lam, P., Alassiri, M. & Lim, T. S. Application of phage display for T-cell receptor discovery. Biotechnol Adv 54, 107870 (2022).

37. Bourgonje, A. R. et al. Phage-display immunoprecipitation sequencing of the antibody epitope repertoire in inflammatory bowel disease reveals distinct antibody signatures. Immunity 56, 1393–1409.e6 (2023).

38. Carrasco, A. et al. Regional Specialisation of T Cell Subsets and Apoptosis in the Human Gut Mucosa: Differences Between Ileum and Colon in Healthy Intestine and Inflammatory Bowel Diseases. J Crohns Colitis 10, 1042–1054 (2016).

39. Van Kruiningen, H. J., West, A. B., Freda, B. J. & Holmes, K. A. Distribution of Peyer’s Patches in the Distal Ileum. Inflamm Bowel Dis 8, 180–185 (2002).

40. Britanova, O. V et al. Targeted depletion of TRBV9+ T cells as immunotherapy in a patient with ankylosing spondylitis. Nat Med 29, 2731–2736 (2023).

41. Kristensen, V. A. et al. Inflammatory bowel disease in South-Eastern Norway III (IBSEN III): a new population-based inception cohort study from South-Eastern Norway. Scand J Gastroenterol 56, 899–905 (2021).

42. Carlson, C. S. et al. Using synthetic templates to design an unbiased multiplex PCR assay. Nat Commun 4, 2680 (2013).

43. Bolotin, D. A. et al. MiXCR: software for comprehensive adaptive immunity profiling. Nat Methods 12, 380–381 (2015).

44. Shannon, P. et al. Cytoscape: a software environment for integrated models of biomolecular interaction networks. Genome Res 13, 2498–2504 (2003).

