## Supplementary figures and tables for "Analyzing the T cell receptor repertoire of 2,804 individuals with inflammatory bowel disease identifies public T cell responses involved in the pathogenesis"

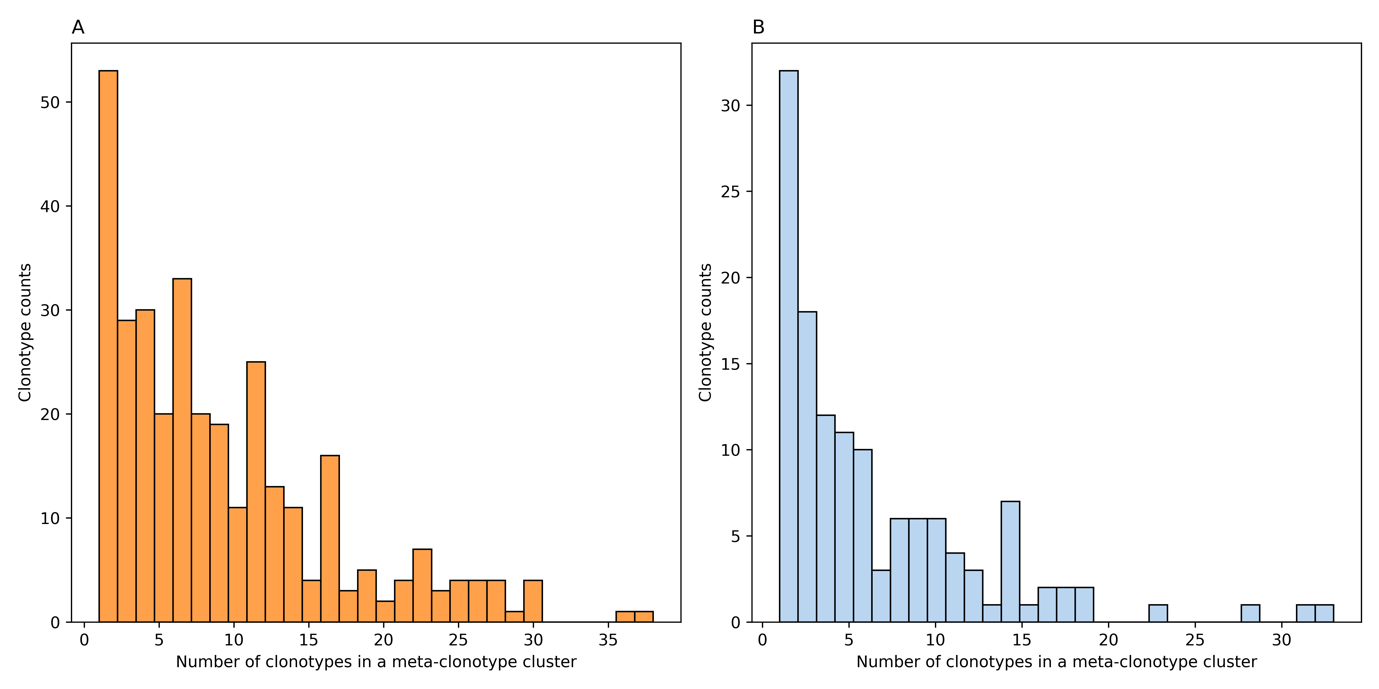

**Figure S1**: The number of clonotypes in each of the 327 and 130 CD- and UC-associated meta-clonotypes. (**A**) depicts the number of clonotypes in each of the 327 CD-associated meta-clonotypes, while (**B**) shows the number of clonotypes in each of the 130 UC-associated meta-clonotypes.

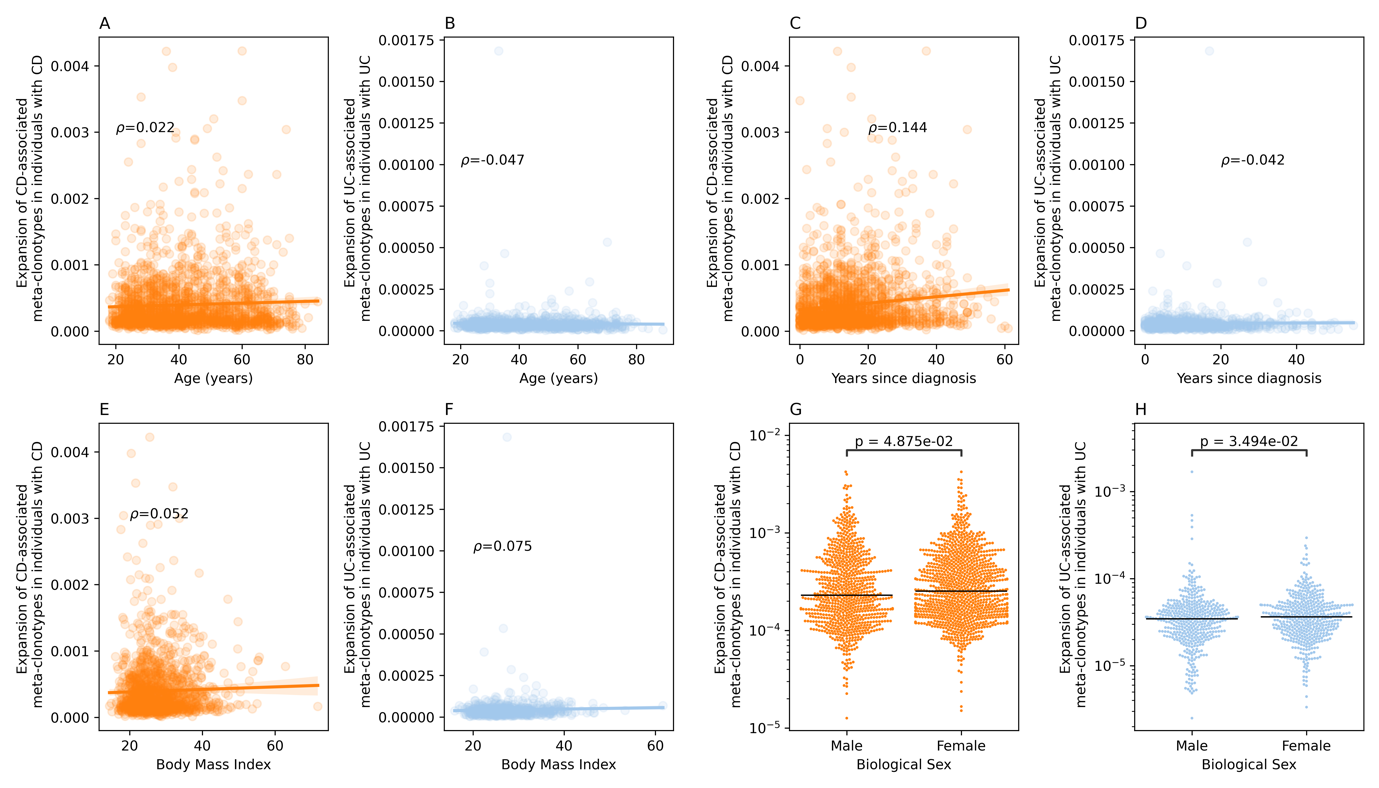

**Figure S2**: The association between the expansion of CD- and UC-associated meta-clonotypes and general phenotypic properties. (**A**) depicts the correlation between the expansion of CD-associated clonotypes and age; the same relationship is shown in (**B**) for UC-associated clonotypes. (**C**) and (**D**) show the correlation between years since diagnosis and the expansion of CD- and UC-associated clonotypes, respectively. (**E**) and (**F**) illustrate the relationship between body mass index (BMI) and the expansion of CD and UC-associated clonotypes, respectively. (**G**) and (**H**) show the expansion of CD- and UC-associated clonotypes in males and females with either CD (**G**) or UC (**H**).

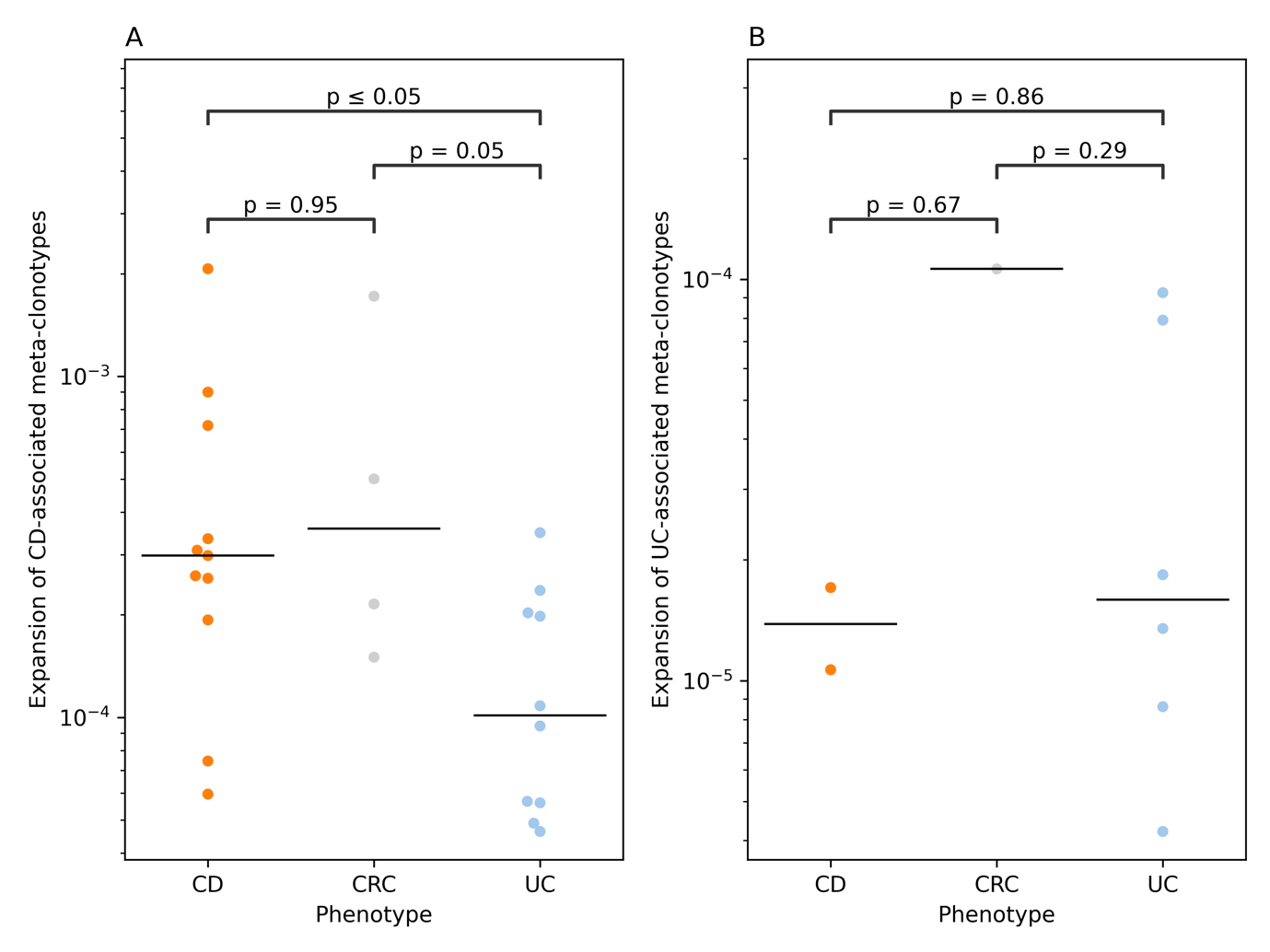

**Figure S3**: The overlap between CD- and UC-associated meta-clonotypes in the bulk surgery samples from Rosati et al.^1^ (**A**) shows the expansion of CD-associated meta-clonotypes in individuals with CD, CRC, or UC. While (**B**) shows the expansion of UC-associated meta-clonotypes in the same phenotypic groups.

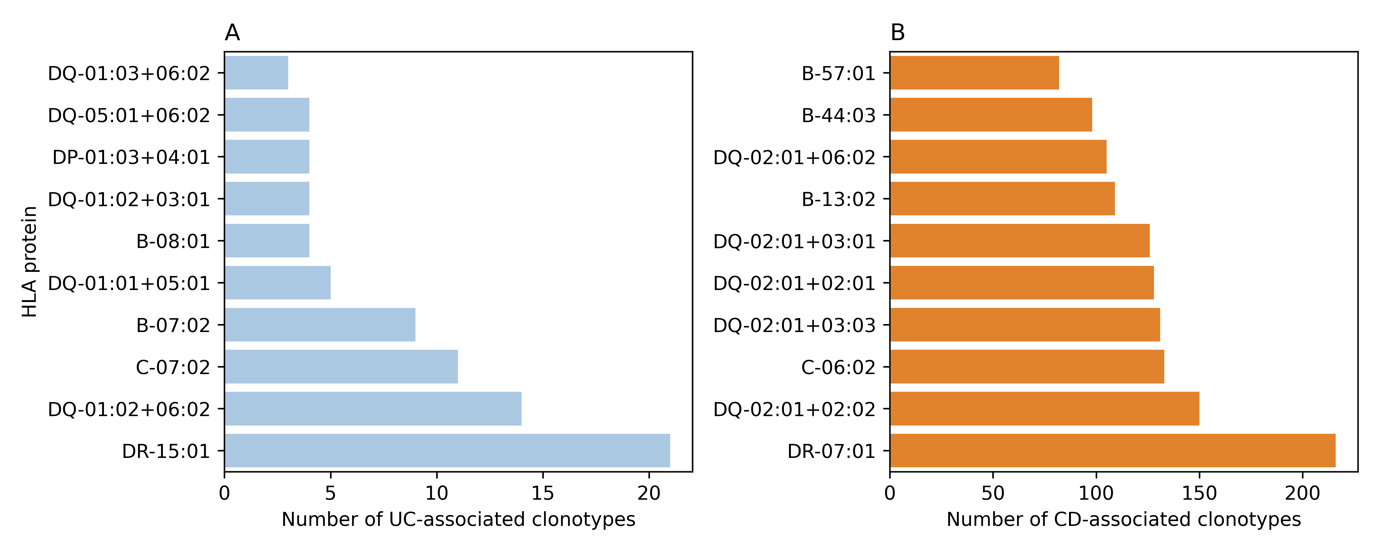

**Figure S4**: The cognate HLA alleles of UC-associated clonotypes (**A**) and CD-associated clonotypes (**B**).
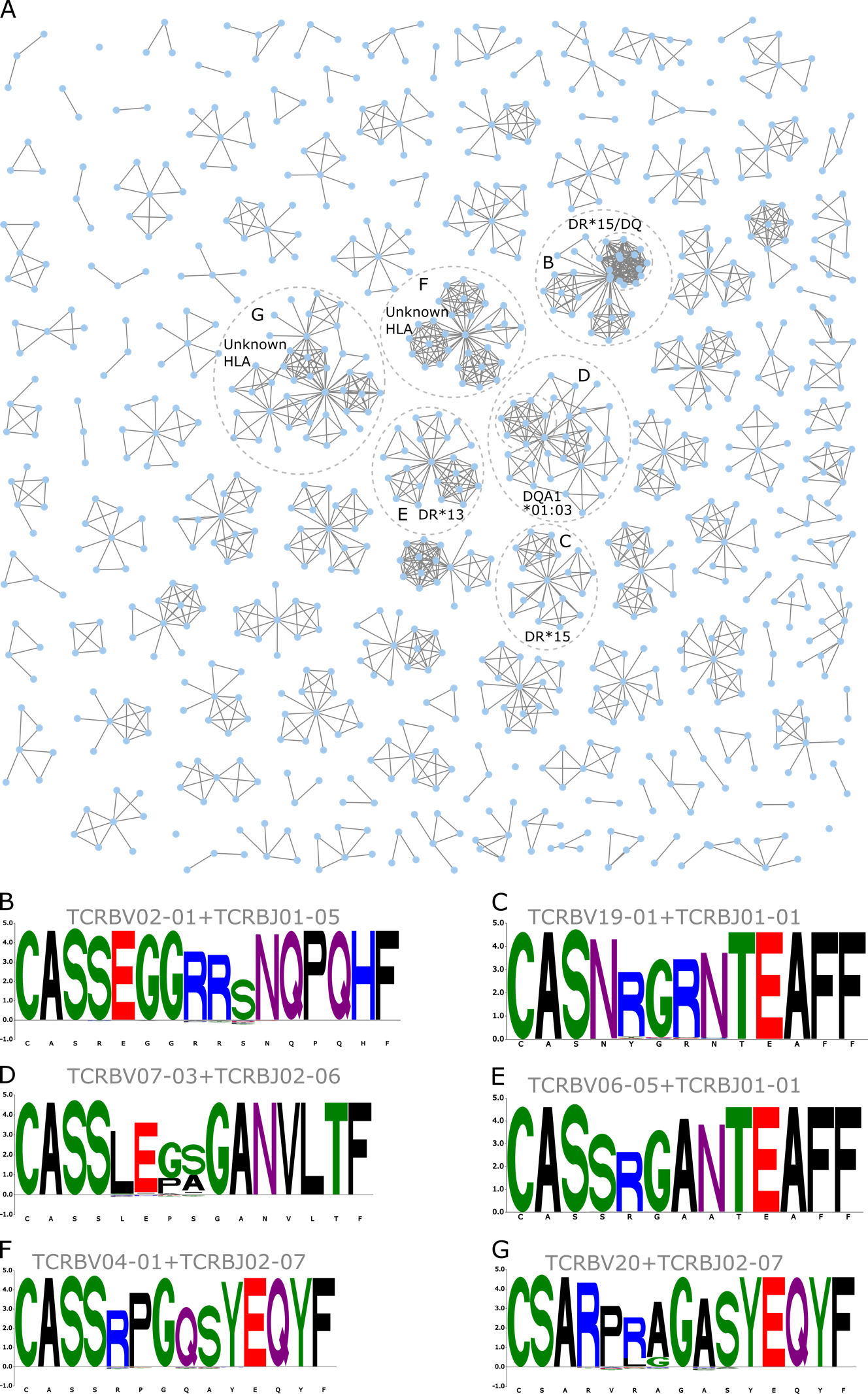

**Figure S5**: Distinct public immune responses associated with UC. (**A**) a graph representation of UC-associated meta-clonotypes, each node in the graph represents a clonotype, and edges represent similarity between these nodes. Specifically, two nodes are connected by an edge if they share the same V and J genes and there is a one-hamming distance between their CDR3 amino acid sequences. (**B**) and (**C**) show the sequence motifs of two sequence clusters that contain multiple clonotypes that are restricted to HLA-DRB1*15:01 alleles or to other alleles on the same haplotype. (**D**) depict the sequence motifs of multiple clonotypes that were restricted to the HLA-DQA1*01:03 alleles, most commonly in conjunction with the HLA-DQB1*05 family. (**E**) depict the sequence motif of a cluster that contains multiple HLA-DRB1*13:01 associated alleles. (**F**) and (**G**) show the sequence motif of two distinct clusters that did not show any association with HLA alleles present in our HLA-TCR association database^2^.

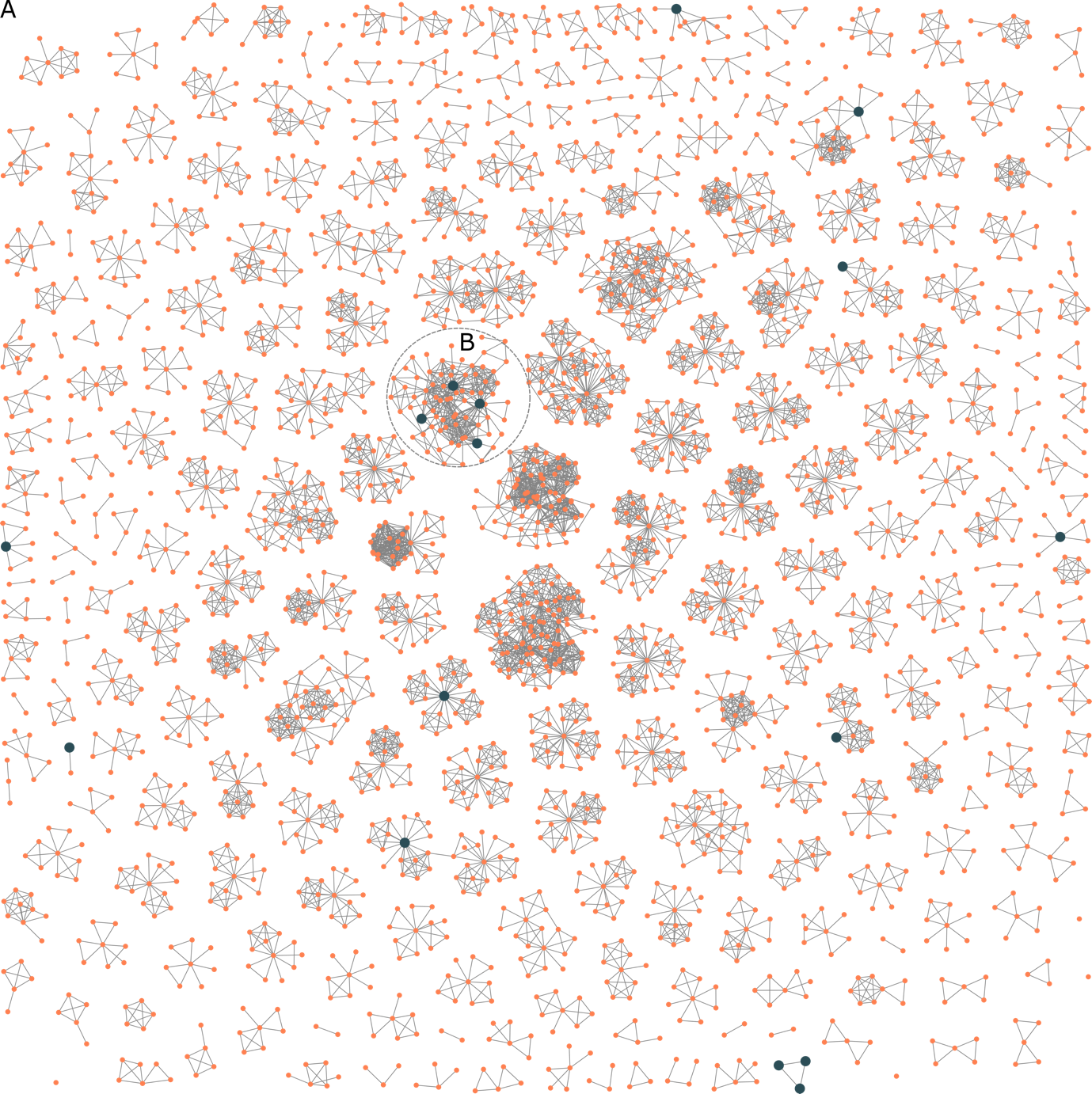

**Figure S6:** The overlap between CD-associated clonotypes and fungal-reactive clonotypes previously described^3^. (**A**) shows a network analysis of CD-associated clonotypes, with orange dots representing CD-associated clonotypes and blue dots representing fungal-reactive clonotypes that are also associated with CD. (**B**) shows the cluster with the most fungal-reactive clonotypes, which belongs to the sequence motif shown in **Fig. 4C**.

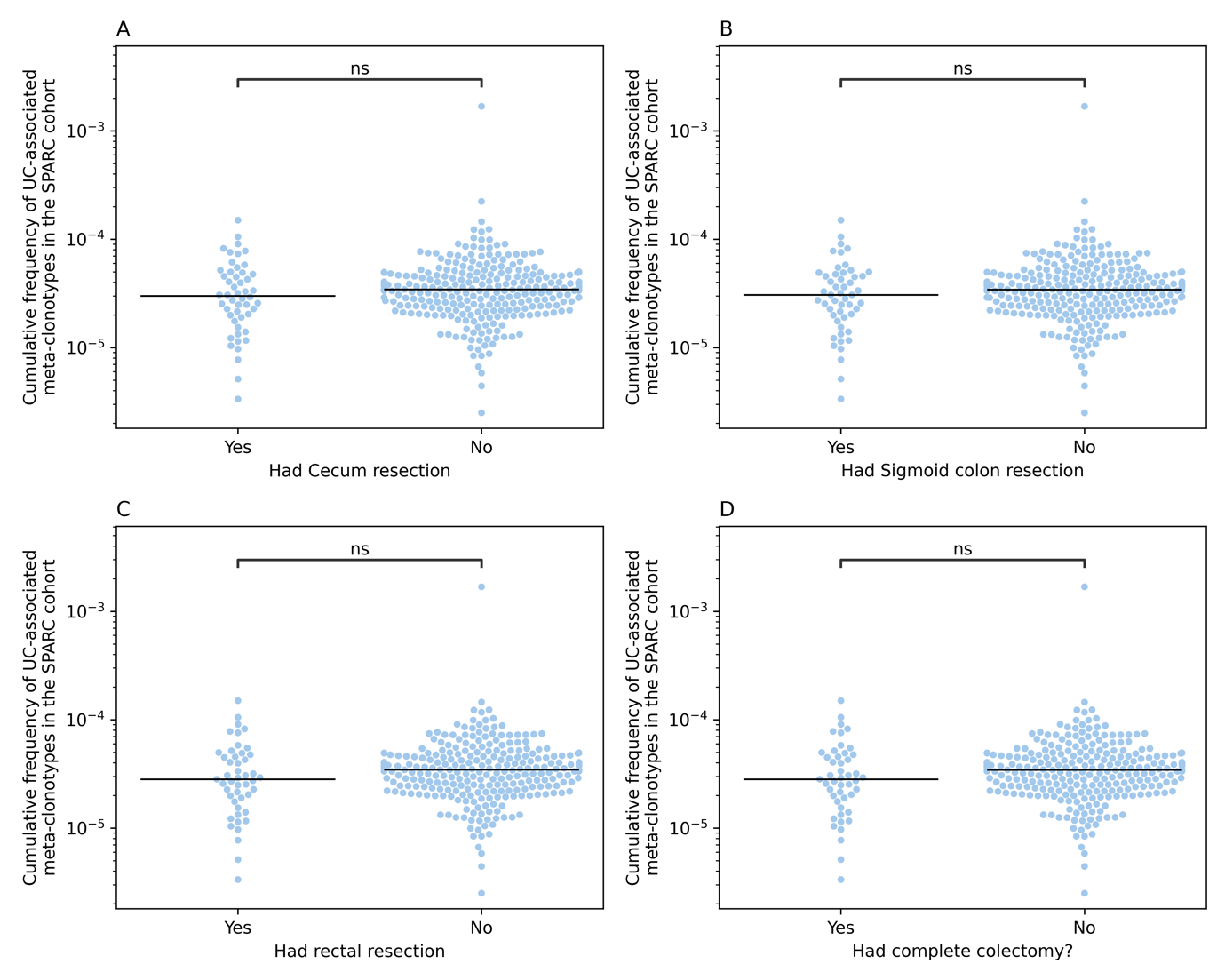

**Figure S7**: The expansion of UC-associated meta-clonotypes in individuals with UC after different surgeries. (**A-D**) show the expansion of UC-associated meta-clonotypes in individuals who underwent resections, namely, cecum (**A**), sigmoid colon (**B**), rectal (**C**), and lastly, complete colectomy (**D**).

**Supplementary Tables**

**Table S1**: Phenotypic properties of the SPARC-cohort.

|  | Crohn’s disease | Ulcerative colitis |
| --- | --- | --- |
| Number of samples | 1,890 | 914 |
| Percentage of females | 56.4% | 50.7% |
| Age at diagnosis | 26.18 ± 12.91 years | 30.97 ± 12.91 years |
| Age at sampling | 41.34 ± 14.3 years | 43.27 ± 15 years |
| Years with disease | 15.13 ± 11.22 years | 12 ± 10.10 years |
| BMI | 27.66 ± 6.63years | 27.44 ± 5.92 |
| Disease location and extension | L1: Ileal (n= 383; 20.2%) | E1: Ulcerative proctitis (n=91; 9.95 %) |
|  | L2: Colonic CD (n= 256; 13.5%) | E2: Left-sided ulcerative colitis (n= 170; 18.59 %) |
|  | L3: Ileocolonic CD (n= 843; 44.6%) | E3: Extensive ulcerative colitis (n=71; **7.76%**) |
|  | Unknown: (n=408; n=21.5%) | E4: Pancolitis (n=389; **42.56%**) |
|  |  | Unknown (193; **21.11%**) |
| Disease BEHAvIOUR | B1: Non‐stricturing, non‐penetrating  (n=742; 39.25 %) | - |
|  | B2: Stricturing (n=417; 22.06 %) | - |
|  | B3: Penetrating (n=253; 13.38 %) |  |
|  | B2+B3: Stricturing and penetrating (n=166; 8.78%) | - |
|  | Unknown (n=312; 16.50%) | - |
| Disease severity  (MAYO-6 category) | - | Mild (n=201; 21.99%) |
|  | - | Moderate (n=86; 9.40 %) |
|  | - | Severe (n=25; 2.73%) |
|  | - | Remission (n=559; 61.15%) |
|  | - | Unknown (n=43; 4.70 %) |
| Disease severity  (SCDAI category) | Mild (n=328; 17.35%) | - |
|  | Moderate (n= 313; 16.5%) | - |
|  | Remission (n= 1,114; 58.9%) | - |
|  | Severe (n= 19; 1%) | - |
|  | Unknown (n=116; 6.3%) | - |

**Table S2:** Phenotypic properties of the IBSEN-III samples included in the study after removing samples with less than 25,000 productive clonotypes.

|  | Crohn’s disease | Ulcerative Colitis | Symptomatic controls |
| --- | --- | --- | --- |
| Total number of samples | 237 | 384 | 240 |
| Number of samples at the treatment-naive state | 215 | 343 | 240 |
| Number of samples at the treated state | 170 | 321 | 0 |
| Number of samples with paired measurements | 148 | 280 | 0 |
| Percentage of females | 53.2% (n = 126) | 44.3% (n = 170) | 49.2% (n = 118) |
| Age at diagnosis | 32 ± 17.4 years | 36.38 ± 14.8 years | 29.1 ± 13.4 years |
| Number of pediatric cases | 64 | 23 | 41 |
| Disease location and extension at the treatment-naive status | L1: Ileal (n= 83; 38.6%) | E1: Ulcerative proctitis (n=133; 38.8%) | **-** |
|  | L2: Colon (n=29; 13.5%) | E2: Left-sided colitis (n=67; 19.5%) | **-** |
|  | L3: Ileocolonic (n=40; 18.6%) | E3: Pancolitis (n=116;33.8%) | **-** |
|  | Unknown (n=63; 29.3%) | Unknown (n=27;7.9%) | **-** |
| Disease modifiers at the treatment-naive status | B1: Non-stricturing, non-penetrating (n=152; 70.7%) | **-** | **-** |
|  | B2: Stricturing (n=30; 13.9%) | **-** | **-** |
|  | B3: Penetrating (n=5; 2.3%) | **-** | **-** |

**Table S3:** The different therapeutic and pharmacological interventions administered to individuals the SPARC IBD cohort.

| Category | Mechanism of Action | Medication | Crohn's disease | Ulcerative colitis |
| --- | --- | --- | --- | --- |
| Biologicals/ monoclonal antibodies | Anti-TNF agents | Infliximab | 46.7% | 34.1% |
|  |  | Adalimumab | 42.7% | 22.5% |
|  |  | Certolizumab pegol | 11.2% | 1.1% |
|  |  | Golimumab | 1.0% | 2.7% |
|  | Anti-integrins | Vedolizumab | 16.6% | 21% |
|  |  | Natalizumab | 1.9% | 0.2% |
|  | Anti-IL12/IL-23 | Ustekinumab | 9% | 2.6% |
| Small molecules | JAK-inhibitors | Tofacitinib | 0.8% | 3.4% |
|  | Anti-proliferative/ Immunosuppressive | Thiopurine | 48.3% | 38.7% |
|  |  | Methotrexate | 20% | 9.4% |
|  | Steroids/ Immunosuppressive | Corticosteroid | 64% | 67% |
|  | anti-inflammatory | Mesalamine | 43.28% | 62.6% |
|  |  | Sulfasalazine | 9.8% | 14.2% |

**Table S4:** *Overview of the different surgeries conducted on the SPARC IBD cohort.*

| Surgery type | Location | Number of surgeries | Crohn's disease | Ulcerative colitis |
| --- | --- | --- | --- | --- |
| All IBD surgery | Any | 0 | 33% | 55% |
|  |  | 1 | 18.5% | 4.1% |
|  |  | 2 | 8.5% | 2.4% |
|  |  | 3 | 4.3% | 1.4% |
|  |  | 4 | 1.7% | 0.1% |
|  |  | 5 | 0.2% | 0% |
|  |  | >5 | 1% | 0.1% |
|  |  | Unknown | 32.8% | 36.8% |
| Small bowel resection | Duodenum | 0 | 52.6% | 45.2% |
|  |  | 1 or more | 0.8% | 0% |
|  |  | Unknown | 46.6% | 54.7% |
|  | Jejunum | 0 | 51.6% | 45.1% |
|  |  | 1 or more | 1.69% | 0.1% |
|  |  | Unknown | 46.76% | 54.8% |
|  | Ileum | 0 | 29% | 42.6% |
|  |  | 1 or more | 26.4% | 2.6% |
|  |  | Unknown | 44.5% | 54.8% |
| Colon resection | Cecum | 0 | 24.5% | 33% |
|  |  | 1 or more | 24.7% | 5.9% |
|  |  | Unknown | 50.6% | 61.1% |
|  | Sigmoid | 0 | 41.2% | 33.6% |
|  |  | 1 or more | 7.6% | 6.12% |
|  |  | Unknown | 51.2% | 60.3% |
|  | Rectum | 0 | 42.8% | 34% |
|  |  | 1 or more | 5.8% | 5.6% |
|  |  | Unknown | 51.4% | 60.4% |
|  | Complete Colectomy | No | 77% | 71% |
|  |  | Yes | 8.3% | 8.5% |
|  |  | Unknown | 14.7% | 20.5% |
